# Biologically informed deep neural network for prostate cancer classification and discovery

**DOI:** 10.1101/2020.12.08.416446

**Authors:** Haitham A. Elmarakeby, Justin Hwang, David Liu, Saud H. AlDubayan, Keyan Salari, Camden Richter, Taylor E Arnoff, Jihye Park, William C. Hahn, Eliezer Van Allen

## Abstract

Determination of molecular features that mediate clinically aggressive phenotypes in prostate cancer (PrCa) remains a major biological and clinical challenge. Here, we developed a biologically informed deep learning model (P-NET) to stratify PrCa patients by treatment resistance state and evaluate molecular drivers of treatment resistance for therapeutic targeting through complete model interpretability. Using a molecular cohort of 1,238 prostate cancers, we demonstrated that P-NET can predict cancer state using molecular data that is superior to other modeling approaches. Moreover, the biological interpretability within P-NET revealed established and novel molecularly altered candidates, such as *MDM4* and *FGFR1*, that were implicated in predicting advanced disease and validated *in vitro*. Broadly, biologically informed fully interpretable neural networks enable preclinical discovery and clinical prediction in prostate cancer and may have general applicability across cancer types.

## Main Text

With the advancement of molecular profiling technologies, the ability to observe millions of genomic, transcriptional, and additional features from cancer patients and their tumors has grown significantly over the past decade. Specifically in prostate cancer, the availability of rich molecular profiling data linked to clinical annotation has enabled discovery of many individual genes, pathways, and complexes that promote lethal castration resistant prostate cancer (CRPC), which has led to both biological investigations and clinical evaluations of these individual features for predictive utility (*1*–*9*). However, the relationships between these molecular features, and their combined predictive and biological contributions to disease progression, drug resistance, and lethal outcomes, remains largely uncharacterized.

When developing a predictive model, one may choose from a large range of approaches, although each comes with tradeoffs of accuracy and interpretability. In translational cancer genomics, interpretability of predictive models is critical, as properties that contribute to the predictive capabilities of the model may not only inform patient care, but also provide insights into the underlying biological processes to prompt functional investigation and therapeutic targeting. Linear models such as logistic regression tend to have high interpretability with less accurate predictive performance, whereas deep learning models often have less interpretability but higher predictive performance (*10*, *11*). Using a typical fully connected dense deep learning approach for building predictive models may also result in overfitting unless the network is well regularized, and such models have a tendency to be computationally expensive and less interpretable (*12*).

Efforts to search for slimmer architecture and sparse networks given a full model demonstrated that sparse models can decrease storage requirements and improve computational performance (*13*–*15*). However, finding such a sparse model may be challenging, since the typical training-pruning-retraining cycle is usually computationally expensive, and recent studies indicate that building a sparse model *de novo* may be easier (*16*). Additionally, efforts to enhance the interpretability of deep learning models and the need to explain model decisions led to the development of multiple attribution methods, including LIME (*17*), DeepLIFT (*10*), DeepExplain (*18*), and SHAP (*19*), that can be used to enhance the deep learning explainability and understand how the model is processing information and making decisions.

Taken together, the advances in sparse model development and attribution methods have informed the development of deep learning models to solve biological problems using customized neural network architectures that are inspired by biological systems. For example, Ma et al, developed a visible neural network system, DCell, to model the effect of gene interaction on cell growth in yeast (*20*). A Pathway-Associated Sparse Deep Neural Network (PASNet) PASNet used a flattened version of pathways to predict patient prognosis in Glioblastoma multiforme (*21*). However, whether biologically informed neural networks can accelerate both biological discovery with translational potential, and simultaneously enable clinical predictive modeling, is largely unknown. Here, we hypothesized that a biologically informed deep learning model built upon advances in sparse deep learning architectures, encoding of biological information, and incorporation of explainability algorithms would achieve superior predictive performance compared to established models and reveal novel patterns of treatment resistance in prostate cancer with translational implications.

## Results

We developed a deep learning predictive model that incorporates prior biologically established hierarchical knowledge in a neural network language to predict cancer state in prostate cancer patients based on their genomic profiles. A set of 3,007 curated biological pathways were used to build a pathway-aware multi-layered hierarchical network (P-NET) (Methods). In P-NET, the patient molecular profile is fed into the model and distributed over a layer of nodes representing a set of genes of interest using weighted links (Figure 1a). Later layers of the network encode a set of pathways with increasing levels of abstraction, whereby lower layers represent fine pathways while later layers represent more complex biological pathways and biological processes. The connections between different layers are constrained to follow known child-parent relationships among encoded features, genes, and pathways, and as a result the network is fully interpretable.

**Fig. 1.**
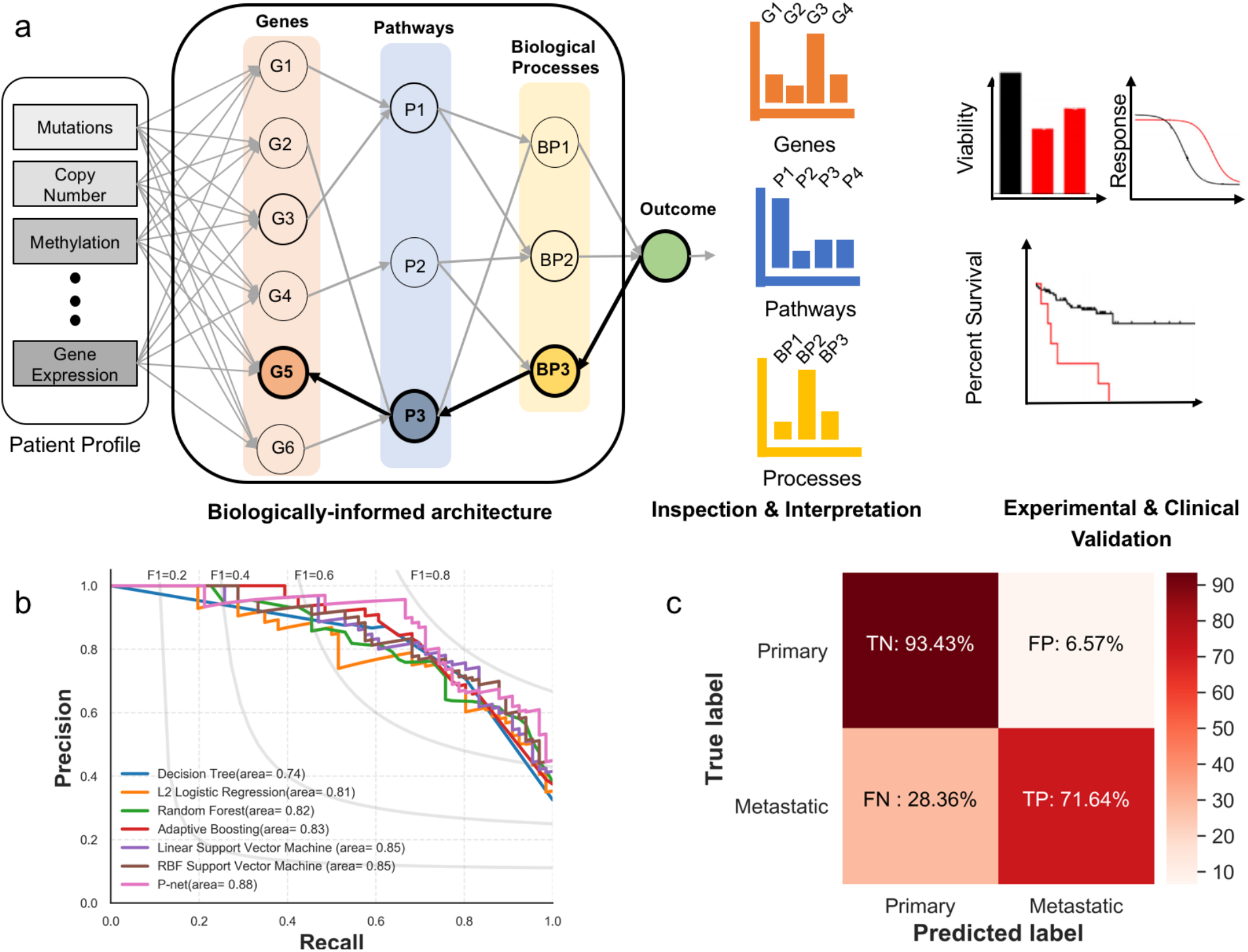
Interpretable biologically informed deep learning model for molecular discovery in metastatic prostate cancer. a) P-NET; neural network architecture that encodes different biological entities into a neural network language with customized connections between consecutive layers (i.e. features from patient profile, genes, pathways, biological processes, and outcome). The trained P-NET provides a relative ranking of nodes in each layer which can be used to generate biological hypotheses regarding the relevance of different biological entities to the outcome of interest. Candidate genes are experimentally and clinically validated to understand the function and the mechanism of action of these genes. b) The Precision-Recall curve of multiple predictive models including Random forest, Support Vector Machine, Decision Trees, and adaptive Boosting as trained and tested on a cohort of prostate cancer patients. P-NET achieves better area under precision recall curve (AUPRC). c) The confusion matrix of P-NET showing the correct classification percentage of samples in the testing set. The model is biased toward predicting more primary samples given that the training set is biased toward having more primary samples (70% primary and 30% metastatic).

We trained and tested P-NET with a set of 1,013 prostate cancers (CRPC = 333, primary = 680), divided into 80% training, 10% validation, and 10% testing, to predict disease state (primary or metastatic disease). The trained P-NET outperformed typical machine learning models, including Support Vector Machine, Logistic Regression, and Decision Trees (AUC=0.93, AUPRC= 0.88, Accuracy 0.83) (Methods, Figure 1b, c, area under ROC curves are shown in Figure S3, 5-fold cross validation results are shown in Figure S4). Furthermore, we evaluated whether the sparse model had characteristics distinct from a dense fully connected deep learning model. We trained a dense model with the same number of parameters as in the P-NET model on training sets with a logarithmically increasing number of samples from 100 to 811 (80% of the total number of samples). The mean performance (determined by AUC) of the P-NET model was higher than the dense model over all the sample sizes, and this difference was statistically significant in smaller sample sizes (<=500) (e.g. mean AUC of 5-fold cross validation was significantly higher for P-NET compared to a dense network trained on 155 samples, *p-value* = 0.003) (Figure 2a, more metrics are shown in Figure S5).

**Fig. 2.**
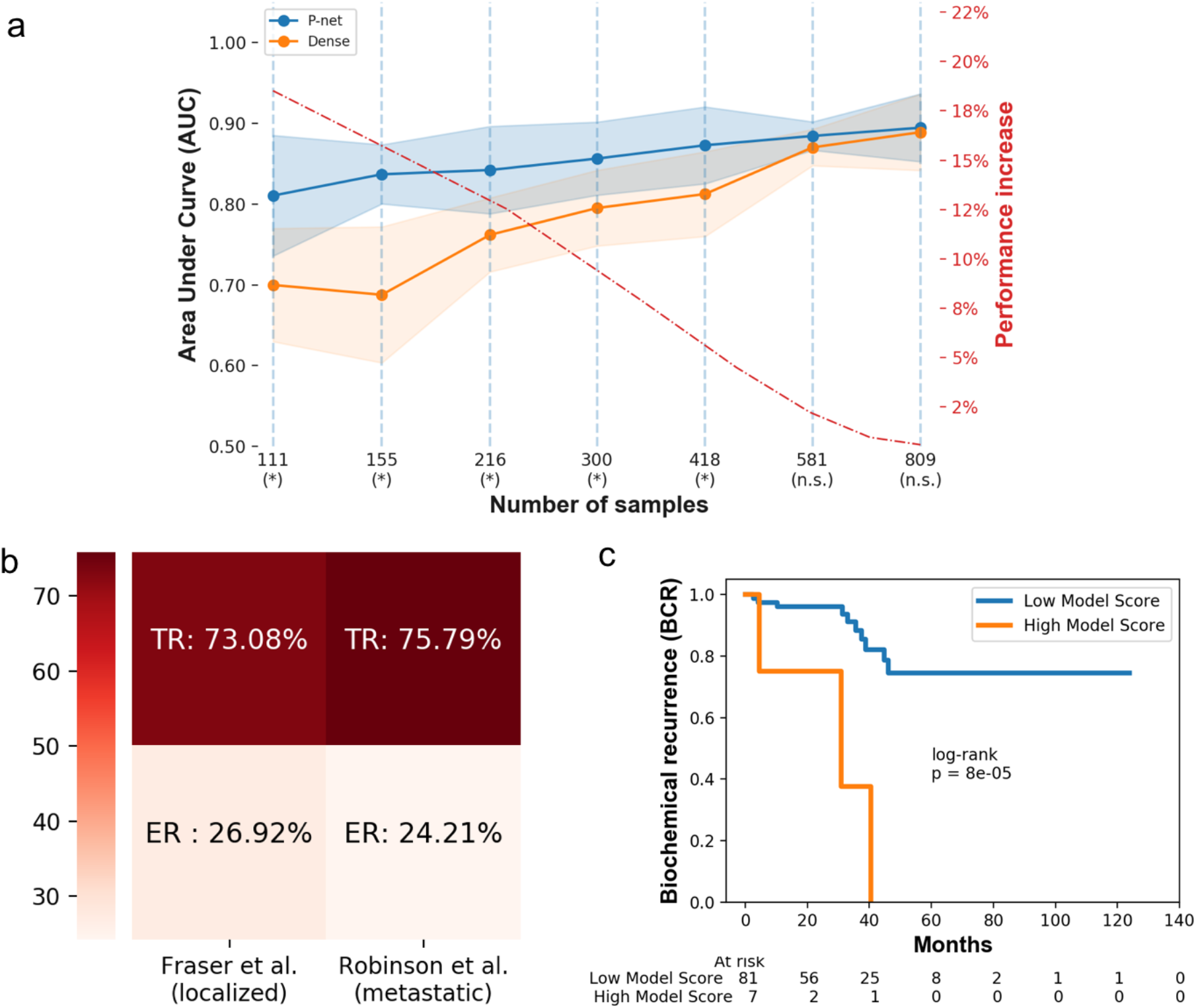
Prediction performance and interpretation of P-NET. a) Prediction performance of P-NET when compared to a fully connected network with the same number of parameters trained using an increasing number of training samples. P-NET achieves better performance (measured as the average AUC over 5 cross validation splits) with smaller numbers of samples compared to a dense fully connected network. The ratio of performance increase is polynomially increasing with the decrease of the number of training samples as shown with the dashed red line. The difference in performance between the two models is statistically significant in all sample sizes less than or equal to 500. Sample sizes marked by (*) indicate statistically significant differences while those marked by (n.s.) are not. b) External validation of P-NET using two independent cohorts (22, 23). The P-NET model achieves 73% and 75.79% true prediction rate (TR) respectively showing that the P-NET can generalize to classify unseen samples. c) Wrong predictions of P-NET were inspected for clinical insights. Patients with high P-NET scores (wrongly classified by P-NET to be resistant samples) have more tendency to have biochemical recurrence (BCR) compared to patients with lower P-NET scores who tend to have progression free survival (PFS). This shows that the P-NET model may be useful in stratifying patients in the clinic and predicting potential BCR before it happens (raw data are included in Data S4).

We next performed external validation of the predictive aspects of the model using two additional PrCa validation cohorts, one primary (*22*) and one metastatic (*23*) (total n=225, links are included in Data S1). The trained P-NET model correctly classified 73% of the primary tumors and 76% of the castration resistant metastatic tumors, indicating that the model can generalize to unseen samples with an adequate predictive performance (Figure 2b). We hypothesized that patients with primary tumors samples incorrectly classified by P-NET as castration resistant metastatic tumors may in fact have worse clinical outcomes. Patients with high P-NET scores misclassified as resistant disease were significantly more likely to have biochemical recurrence than patients with low P-NET scores (*p* = 8*10-5; log-rank) indicating that for patients with primary prostate cancer, the P-NET score may be used to predict potential biochemical recurrence (Figure 2c).

To understand the interactions between different features, genes, pathways, and biological processes that contributed to the predictive performance, and to study the paths of impact from the input to the outcome, we visualized the whole structure of P-NET with the fully interpretable layers after training (Figure 3a and S6). Among aggregate molecular alterations, copy number variation (CNV) was more informative compared to mutations, consistent with prior reports (*24*). In addition, P-NET selected a hierarchy of pathways (out of 3,007 pathways on which P-NET was trained) as relevant to classification, including post-translational modification (PTM) (including Ubiquitination and SUMoylation) and transcriptional regulation by *RUNX2* and *TP53*. Ubiquitination and SUMoylation pathways contribute to the regulation of multiple tumor suppressors and oncogenes, and dysregulation of these pathways has been linked to prostate cancer initiation and progression in preclinical models (*25*). *RUNX2* is an osteogenic transcription factor that regulates cell proliferation and is associated with metastatic disease in prostate cancer patients (*26*).

**Fig. 3.**
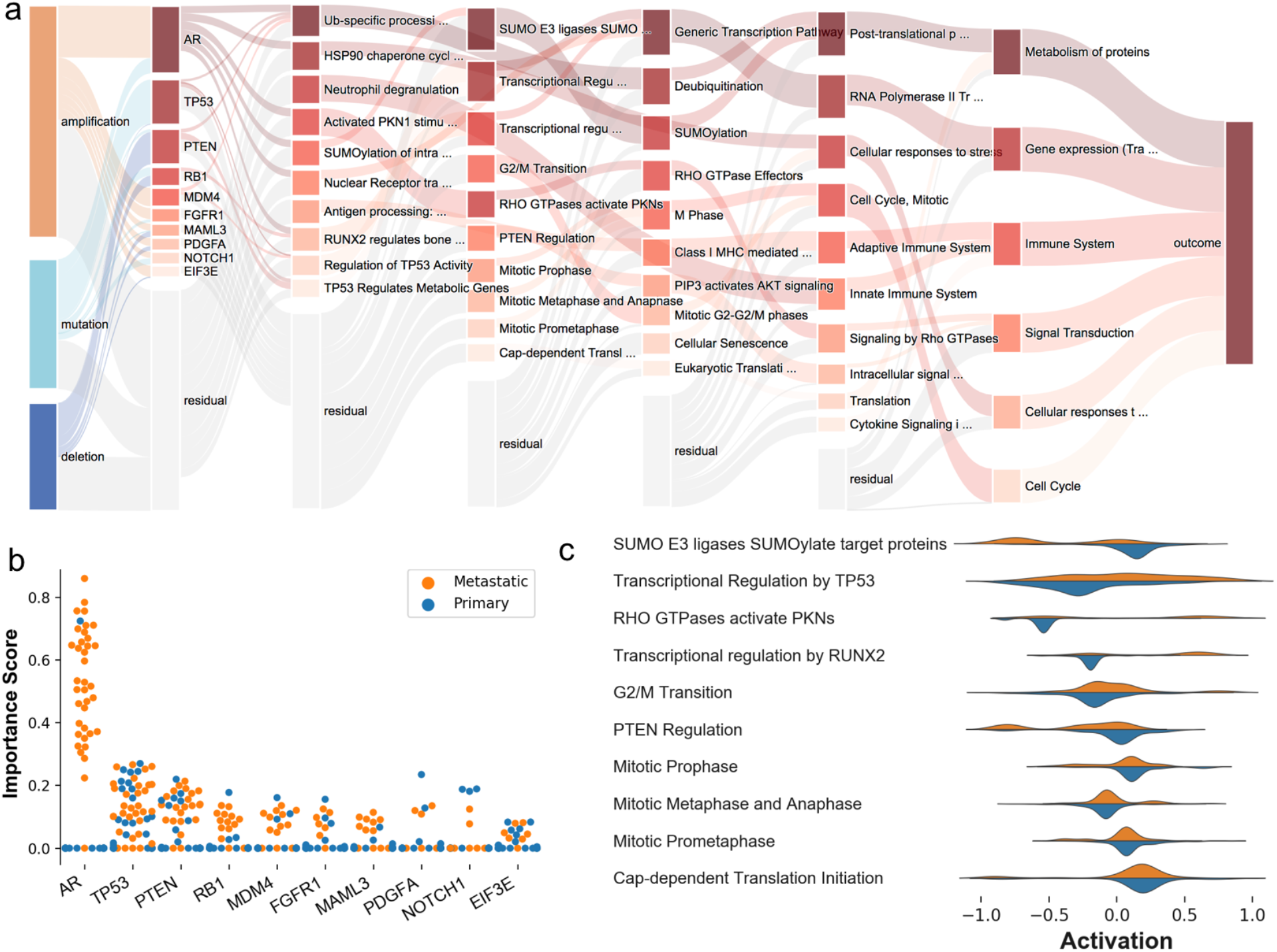
Inspecting and interpreting P-NET. a) Visualization of inner layers of P-NET shows the estimated relative importance of different nodes in each layer. Nodes on the far left represent feature types (mutation and copy number amplification and deletion). The second layer nodes represent genes and layers representing higher level biological entities shown in layers to the right. The final layer represents the model outcome. Nodes with darker color are shown to be more important while nodes with transparent color represent the residual importance of non-shown nodes in each layer. The contribution of datatype to the importance of each gene is depicted using the Sankey diagram. The importance of *AR* gene is driven mainly by gene amplification, the importance of *TP53* is driven by mutation, and the importance of *PTEN* gene is driven by deletion. b) Top genes are ranked based on the average importance of each gene. The distribution of sample-level importance calculated for the testing set is shown in the Swarm diagram. In general, metastatic samples tend to have more influence on the importance of different genes. c) Inspecting the activation output of each node shows how the outcome of different nodes changes with changing the class of input samples (primary-blue vs. metastatic-orange). The activation distribution of top nodes in layer 3 shows that the “Transcription Regulation by *TP53*” pathways is ranked second among pathways in the same layer. Metastatic samples tend to push the activation of this pathway toward positive values while primary samples tend to have negative activation of this pathway.

To evaluate the relative importance of specific genes contributing to the model prediction, we inspected the genes layer and utilized the DeepLIFT attribution method to obtain the average ranking of genes (Methods) (*10*). Highly ranked genes included *AR*, *PTEN*, *RB1*, and *TP53*, which are known PrCa drivers previously associated with metastatic disease (*2*, *5*, *7*, *27*). In addition, alterations in less expected genes, such as *MDM4*, *FGFR1, NOTCH1,* and *PDGFA*, strongly contributed to predictive performance (Figure 3b). To understand the behavior of trained P-NET, we checked the activation of each node in the network and asked whether this activation changed with the change of the input sample class (Primary vs. Metastatic) (Methods). We observed that the difference in the node activation was higher in higher layers and more concentrated in highly ranked nodes in each layer (Figure S7). For example, the activation distribution of the nodes of layer 3 was different when P-NET was given a primary sample compared to a resistant sample (Figure 3c). Thus, the interpretable architecture of P-NET can be interrogated to understand how the input information is transformed through layers and nodes, enabling further understanding of the state and importance of the involved biological entities.

Through evaluation of multiple layers in the P-NET trained model, we observed convergence in *TP53-*associated biology contributing to CRPC. Tracing the relevance of *TP53*-related pathways to the gene levels, we detected that in addition to *TP53* and *MDM2*, each established in prostate cancer disease progression (*27*–*31*), alterations in *MDM4* strongly contributed to this network convergence. *MDM4* can inhibit wild-type *TP53* expression by binding to and masking the transcriptional activation domain, though its role in prostate cancer treatment resistance is incompletely characterized (*32*).

We further studied the *MDM4* profile both in clinical samples and functional models. *MDM4* focal amplification was more prevalent in resistant samples compared to primary samples (X2Yates correction = 40.8251, *p-value* <0.00001) (Figure 4a). Copy number and mutation distributions of other top genes are shown in Figure S8. In a genome-wide gain-of-function preclinical screen using 17,255 open reading frames ORFs in LNCaP cells, *MDM4* overexpression was significantly associated with resistance to enzalutamide, a second-generation antiandrogen medication which is used in CRPC patient populations (*33*) (Figure 4c). We thus utilized CRISPR-Cas9 to target *MDM4* in LNCaP prostate cancer cell lines. In comparison to a negative control, proliferation of LNCaP cells was reduced by 60% (*p-value* <0.0001) (Figure 4d) in response to *MDM4* depletion using two distinct sgRNAs (Figure 4e). This indicated that selective therapeutic targeting of *MDM4* may be viable in TP53-wildtype advanced prostate cancer patients. We then sought to study the effect of inhibiting *MDM4* in prostate cell lines with mutant and wild-type *TP53*. Prostate cells with wild-type TP53 were more sensitive to the *MDM4* inhibitor RO-5963 compared to mutant *TP53* cell lines (2μM of RO-5963 reduced the viability of LNCaP cells by nearly 50%; Figure 4f). Overall, convergence of TP53 pathway dysregulation across multiple layers of the trained P-NET model identified specific vulnerabilities involving *MDM4*, which can be therapeutically targeted with *MDM4*-selective inhibition in a genomically stratified prostate cancer patient population.

**Fig. 4.**
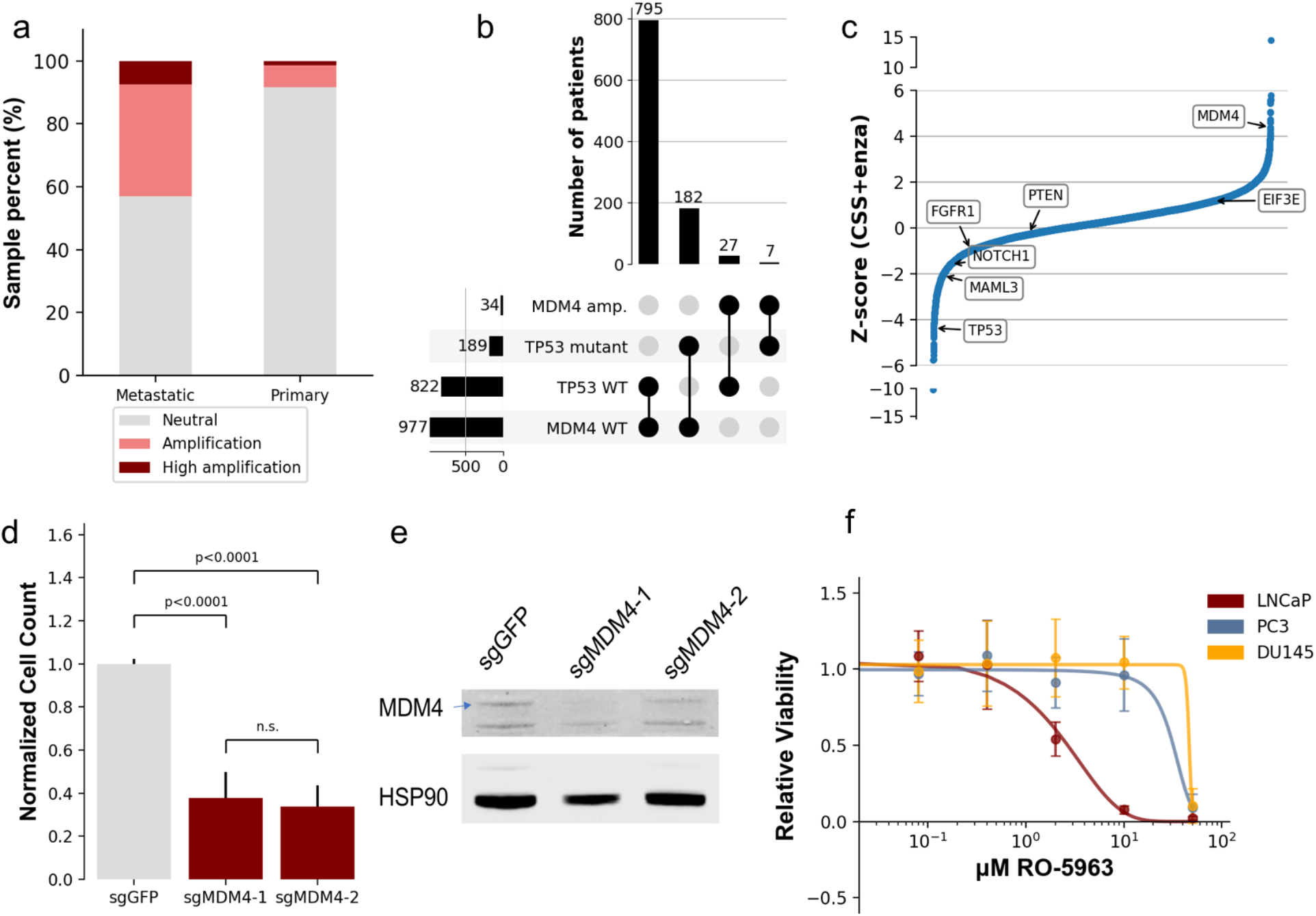
Functional validation. a) Distribution of *MDM4* alterations across 1,013 PrCa samples showing more prevalence of *MDM4* amplification in resistant samples compared to primary samples. b) Joint distribution of TP53 mutations and *MDM4* amplifications across 1,013 PrCa samples. c) Analysis of enzalutamide resistant genes in LNCaP cells based on a genome-scale screen including 17,255 ORFs. The relative enzalutamide resistance of each ORF (x-axis) is plotted as a Z-score (y-axis) (raw data are included in Data S2). A positive Z-score indicates that the gene promotes resistance. *MDM4* and other hits are highlighted on the graph, with *MDM4* scoring as the strongest hit. D) Relative viability of LNCaP cells after transduction of CRISPR-Cas9 and sgRNAs targeting *MDM4* (2 guides, red) or control GFP (black). Data represents the mean + SD of seven replicates (raw data are included in Data S3). e) Immunoblot confirming MDM4 gene deletion in LNCaP cells. HSP90 is a loading control. f) Sensitivity of LNCaP, PC3, and DU145 cells to RO-5963. Relative viability is shown at each indicated dosage of RO-5963. Data represents the mean + SD of six replicates (raw data are included in Data S3)

## Discussion

Broadly, P-NET leveraged a biologically informed, rather than arbitrarily overparameterized, architecture for prediction. As a result, P-NET dramatically reduced the number of parameters for learning, which led to enhanced interpretability. The sparse architecture in P-NET has better predictive performance when compared to other machine learning models, including dense networks, and may be applicable to other similar tasks. Application of P-NET to a molecular cohort of prostate cancer patients demonstrated (i) model performance that may enable prediction of clinically aggressive disease in primary prostate cancer patient populations, and (ii) convergent biological processes that contribute to a metastatic prostate cancer clinical phenotype that harbor novel therapeutic strategies in molecularly stratified populations.

Furthermore, P-NET provided a natural way for integrating multiple molecular features (e.g. mutations and copy number variations) weighted differently to reflect their importance in predicting the final outcome, which previously required different significance approaches for each feature to enable cancer gene discovery (*34*, *35*). Even more, P-NET provided a framework for encoding hierarchical prior knowledge using neural network languages, and turning these hierarchies into a computational model that can be used both for prediction and biological discovery. Specifically, P-NET accurately predicted advanced prostate disease based on the patient's genomic profile and had the ability to predict potential BCR before happening. P-NET visualization enabled a multilevel view of the involved biological pathways and processes, which may guide researchers to develop hypotheses regarding the underlying biological processes involved in cancer progression and translate these discoveries into therapeutic opportunities. Specifically, P-NET rediscovered known genes such as *AR, PTEN, TP53*, and *RB1*. Moreover, P-NET nominated *MDM4* as a novel therapeutic target, which was experimentally validated and may inform further genomically directed therapeutic options for this patient population.

While P-NET provides a framework for outcome prediction and hypothesis generation, this model still requires tuning and training before being used in any given context. As with all deep learning models, the final trained model heavily depends on the hyperparameters used to train the model. In addition, P-NET encodes biological pathways inside the network in a hardcoded way, which makes the model depend on the quality of the annotations used to build the model. Thus, the portability of this approach across different histological and clinical contexts requires further evaluation.

In conclusion, P-NET, a biologically informed deep neural network, accurately classified castration resistant metastatic versus primary prostate cancers. Visualizing the trained model generated novel hypotheses for mechanisms of metastasis in prostate cancer, and provided insights with direct potential for clinical translation in molecularly stratified prostate cancer patient populations. Biologically guided neural networks represent a novel approach to integrating cancer biology with machine-learning by building mechanistic predictive models, providing a platform for biological discovery that may be broadly applicable across cancer prediction and discovery tasks

## Author contributions

Conceptualization, H.A.E. and E.M.V.; Methodology, H.A.E. and E.M.V.; Software, H.A.E.; Formal analysis, H.A.E., D.L., S.H.A., K.S., J.P., and E.M.V.; Investigation, J.H., C.R., T.E.A, H.A.E., W.C.H., and E.M.V.; Writing – Original Draft, H.A.E., J.H., C.R., T.E.A, and E.M.V.; Writing – Review & Editing, H.A.E., J.H., D.L., S.H.A., K.S., C.R., T.E.A, J.P., W.C.H., and E.M.V.; Visualization, H.A.E., J.H., C.R., T.E.A, and E.M.V.; Supervision: W.C.H. and E.M.V.; Funding Acquisition, H.A.E., W.C.H., and E.M.V.

## Data and materials availability

All data is available in the main text or the supplementary materials.

## Methods

We introduce P-NET, an artificial neural network with biologically informed, parsimonious architecture that accurately predicts metastasis in PrCa patients based on their genomic profiles. P-NET is a feedforward neural network with constraints on the nodes and edges. In P-NET, each node encodes some biological entity (e.g. genes and pathways) and each edge represents a known relationship between the corresponding entities. The constraints on the nodes allow for better understanding of the state of different biological components. The constraints on the edges allow us to use a large number of nodes without increasing the number of edges, which leads to a smaller number of parameters compared to fully connected networks with the same number of nodes, and hence potentially less computations. The architecture was built using the Reactome pathway data sets (*36*). The whole Reactome dataset was downloaded and processed to form a layered network of five layers of pathways, one layer of genes, and one layer for features. This sparse model had slightly over 71,000 weights with the number of nodes per layer distributed as shown in Fig S1.B. A dense network with the same number of nodes would have more than 270 million weights with the first layer containing more than 94% of the weights. A hybrid model which contains a sparse layer followed by dense layers still contains over 14 million weights. The number of dense weights is calculated as *n*_*l*_ = (*n*_*l*-1_ + 1) where *w*_*l*_ is the number of weights per layer *l* and *n*_*l*_ is the number of nodes of the same layer.

The meaning of the nodes, layers, and connection of P-NET is encoded through a carefully engineered architecture and a set of restrictions on the connections of the network. The input layer is meant to represent features that can be measured and fed into the network. The second layer represents a set of genes of interest. The higher layers represent a hierarchy of pathways and biological processes that are manually curated. The first layer of P-NET is connected to the next layer via a set of one-to-one connections where each node in the next layer is connected to exactly three nodes of the input layer representing mutations, copy number amplification, and copy number deletions. This scheme results in a much smaller number of weights in the first layer compared to a fully connected network and the special pattern of the connection matrix results in more efficient training. The second layer is restricted to have connections reflecting the gene-pathway relationships as curated by the Reactome pathway dataset. The connections are encoded by a mask matrix *M* that is multiplied by the weights matrix *W* to zero-out all the connections that don't exist in the Reactome pathway dataset. For the next layers, a similar scheme is devised to control the connection between consecutive layers to reflect the real parent-child relationships that exist in the Reactome dataset. The output of each layer is calculated as *y* = *f*[(*M* * *W*) *x* + *b*] where *f* is the activation function, *M* is the mask matrix, *W* is the weights matrix, *x* is the input matrix, and *b* is the bias vector (see Fig. S1.A). The activation of each node is kept into the range of [−1,1] by applying the tanh function *f* = *tanh* = (*e*^2*x*^ − 1)/ (*e*^2*x*^ + 1) to the weighted inputs of the node. The activation of the outcome layers is calculated by the sigmoid function *σ* = 1/(1 + *e*^-*x*^).

To allow each layer to be useful by itself, we added a predictive layer after each hidden layer. P-NET has a smaller number of nodes per layer in the later layers compared to the first layers Fig. S1.B. Since it is more challenging to fit the data using a smaller number of weights in the later layers, we used a higher loss weight for later layer outcomes during the optimization process. The final prediction of the network was calculated by taking the average of all the layer outcomes, Fig S1.C. The learning rate was initialized to be 0.001 and actively reduced after every 50 epochs to allow for smooth convergence. Since we have an unbalanced dataset, we weighted the classes differently to reduce the network bias toward one class based on the bias in the training set. The model was trained using Adam optimizer (*37*) to reduce the binary cross-entropy loss functions 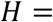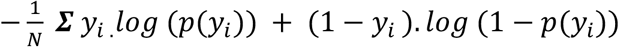 is the probability that sample *i* has a metastatic cancer as calculated using the sigmoid function σ, and *N* is the total number of samples. We checked different gradient based attribution methods to rank the features in all the layers, and we chose to use the DeepLIFT scheme as implemented in the DeepExplain library (*10*).

To check the utility of the developed model, we trained P-NET to predict cancer state (primary/metastatic) of prostate patients based on their genomic profiles. We used whole-exome sequencing of 1,013 patients along with the corresponding somatic mutations and copy number alterations (*4*). The mutations were aggregated on the gene level, excluding Silent, Intron, 3'UTR, 5'UTR, RNA, and lincRNA mutations (Fig. S2). The copy number alterations for each gene was assigned based on the called segment level copy number emphasizing focal amplifications and deletions and excluding single copy amplification and deletions. The prediction performance was measured using the average area under the ROC curve (AUC), the area under precision-recall curve (AUPRC), and F1 score. The corresponding measures were reported for the testing split and also for the cross-validation setup. The input data was divided into a testing set (10%) and a development set (90%). The development set was further divided into a validation set that has the same size as the testing set and the remaining samples are reserved for training. For the cross-validation experiments, the development data set was divided into 5 folds stratified by the label classes to account for the bias in the dataset. The implementation of the proposed system along with the reproducible results are available on Github (https://github.com/marakeby/pnet_prostate_paper).

### Analysis of a genome-scale ORF screen

A genome-scale ORF screen was previously performed in LNCaP cells (33). In brief, cells were infected with a pooled ORF library, subject to puromycin selection to isolate cells containing the respective ORFs, and then seeded in low androgen media (CSS) with enzalutamide. The relative effect of each ORF on cell proliferation was determined after 25 days in culture and is represented as Z-scores. We postulated that amplified genes identified by P-NET regulate oncogenic functions in metastatic castration resistant prostate cancer. To validate this hypothesis, we analyzed this previously published genome-scale ORF screen performed in LNCaP cells which identified genes that, when overexpressed, promoted resistance to the AR inhibitor, enzalutamide (Figure 4C) (*33*). LNCaP cells are dependent on AR and treatment with enzalutamide attenuates cell proliferation. Based on this analysis, *MDM4* scored as a robust enzalutamide resistant gene relative to other hits, including cell cycle regulators (*CDK4, CDK6*) or those with roles in FGF signaling (*FGFR2, FGFR3, FGF6*); these are two pathways implicated in driving resistance to anti-androgen therapies in clinical prostate cancers (*38*, *39*).

### Sensitivity to RO-5963

LNCaP, DU145, and PC3 cells were seeded in 12-well plates at 100k, 20k, or 20k, respectively. After 24 hours, cells were treated with increasing concentrations of RO-5963 between 80nM and 50μM. Media containing the inhibitor was refreshed after 3 days. Relative cell viability was determined using a Vi-Cell after 6 days of treatment, and cell counts were used to calculate IC50 values.

### *MDM4* gene depletion experiments

Blasticidin-resistant Cas9 positive LNCaP cells were cultured in 150μg/mL blasticidin (Thermo Fisher Scientific, NC9016621) for 72 hours to enrich cells with optimal Cas9 activity. 2 million cells were seeded in parallel 10cm plates and infected with lentiviruses expressing puromycin-resistant sgRNAs targeting *MDM4* or GFP control 24 hours later. Cells were then subject to puromycin selection for 4 days, at which point one plate was harvested for immunoblotting and the other was counted using a Vi-Cell and seeded for a proliferation assay. 7 days later, cells were counted again with a Vi-Cell to assess viability, representing a total of 12 days. The target sequence against GFP was CACCGGCCACAAGTTCAGCGTGTCG (sgGFP). The target sequences against *MDM4* were AGATGTTGAACACTGAGCAG (sgMDM4-1) and CTCTCCTGGACAAATCAATC (sgMDM4-2).

### Immunoblotting

Cell pellets were lysed in RIPA buffer (MilliporeSigma, 20-188) containing Protease/Phosphatase Inhibitor Cocktail (Cell Signaling Technology, 5872S). Protein concentrations were calculated using a Pierce BCA Protein Assay Kit (Thermo Fisher Scientific, PI23225), and protein was then denatured in NuPAGE LDS sample buffer (Thermo Fisher Scientific, NP0007) with 5% β-Mercaptoethanol. 13μg of each protein sample was electrophoresed using NuPAGE 4-12% Bis-Tris Protein gels (Thermo Fisher Scientific) and run with NuPAGE MOPS SDS Running Buffer (Thermo Fisher Scientific, NP0001). Proteins were transferred to nitrocellulose membranes using an iBlot apparatus (Thermo Fisher Scientific). Membranes were blocked in Odyssey Blocking Buffer (LI-COR Biosciences, 927-70010) for one hour at room temperature, and membranes were then cut and incubated in primary antibodies diluted 1:1000 in Odyssey Blocking Buffer at 4°C overnight. The following morning, membranes were washed with Phosphate-Buffer Saline, 0.1% Tween (PBST) and incubated with fluorescent anti-rabbit secondary antibodies (Thermo Fisher Scientific, NC9401842) for one hour at room temperature. Membranes were again washed with PBST and then imaged using an Odyssey Imaging System (LI-COR Biosciences). Primary antibodies used include MDM4 (Thermo Fisher Scientific, A300287A) and HSP90 (Cell Signaling Technology, 4874S).

### Chemical inhibition of MDM4 reduces prostate cancer cell viability

Given the proposed role that *MDM4* plays in driving enzalutamide resistance in prostate cancer cells, we sought to determine the response of prostate cancer cells to chemical inhibition of *MDM4*. We evaluated RO-5963, a small molecule *MDM2/4* dual inhibitor with the greatest selectivity towards MDM4 in its class (*40*). This drug has previously demonstrated robust efficacy against *MDM4* dependent cancer cell lines (*41*). We evaluated the effects of increasing concentrations of RO-5963 on prostate cancer cell proliferation.

### Gene depletion of *MDM4* reduces prostate cancer cell viability

To determine how prostate cancer cells would respond to precision tools that target *MDM4* at the gene level, we utilized CRISPR-Cas9 and two sgRNAs targeting distinct sequences of *MDM4* in LNCaP cells. In comparison to a negative control sgRNA (GFP), viability of LNCaP cells was reduced by about 60% (Figure 4D) in response to *MDM4* depletion (Figure 4F) after 12 days in culture. Altogether, we concluded that *MDM4* regulates enzalutamide resistance, and that targeting *MDM4* through either chemical or genetic approaches significantly attenuated the viability of prostate cancer cell lines. Our observations indicate that antagonizing *MDM4* in metastatic castration resistant prostate cancers that harbor wild-type *p53* is an attractive precision strategy.

**Fig. S1.**
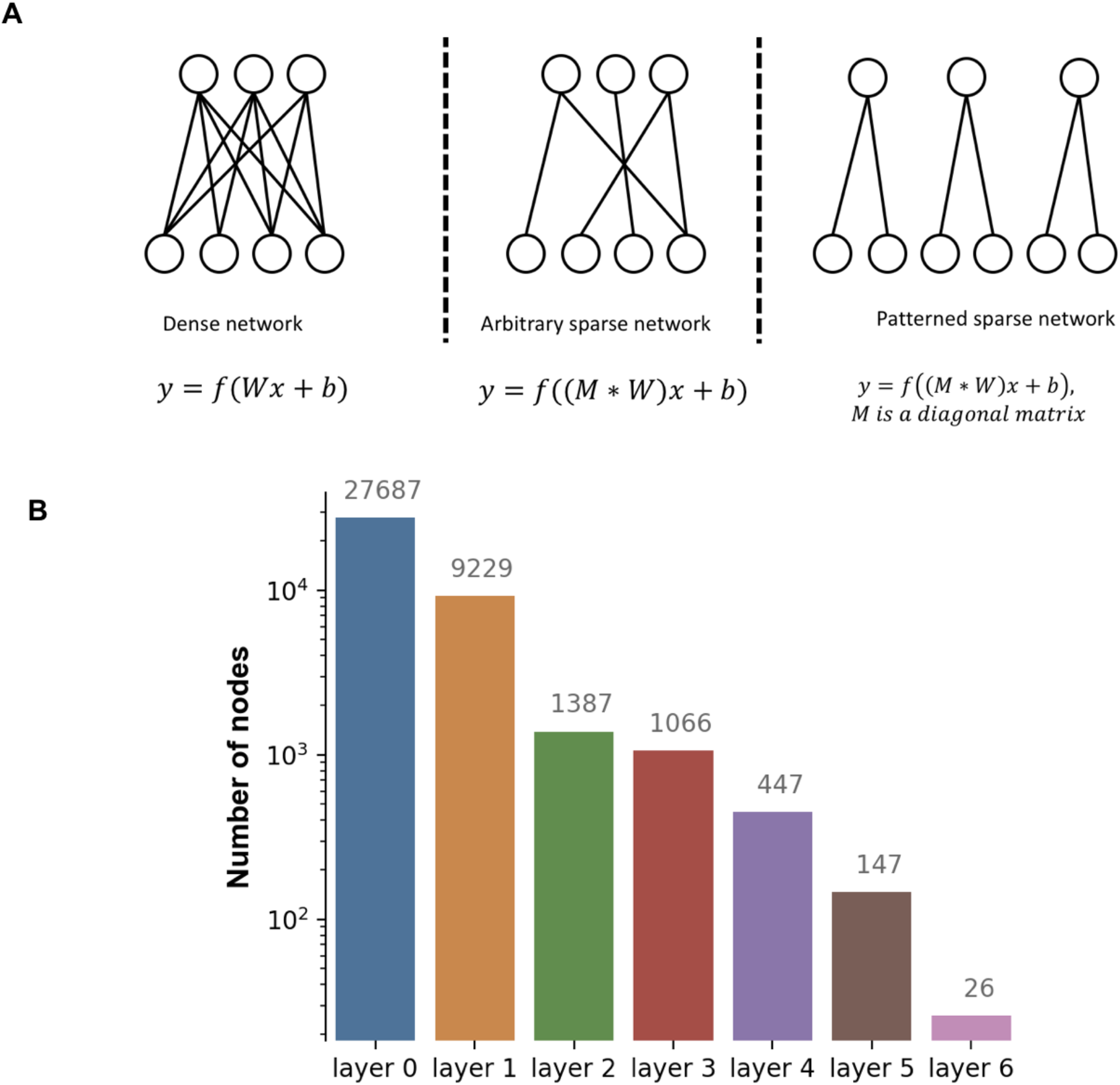
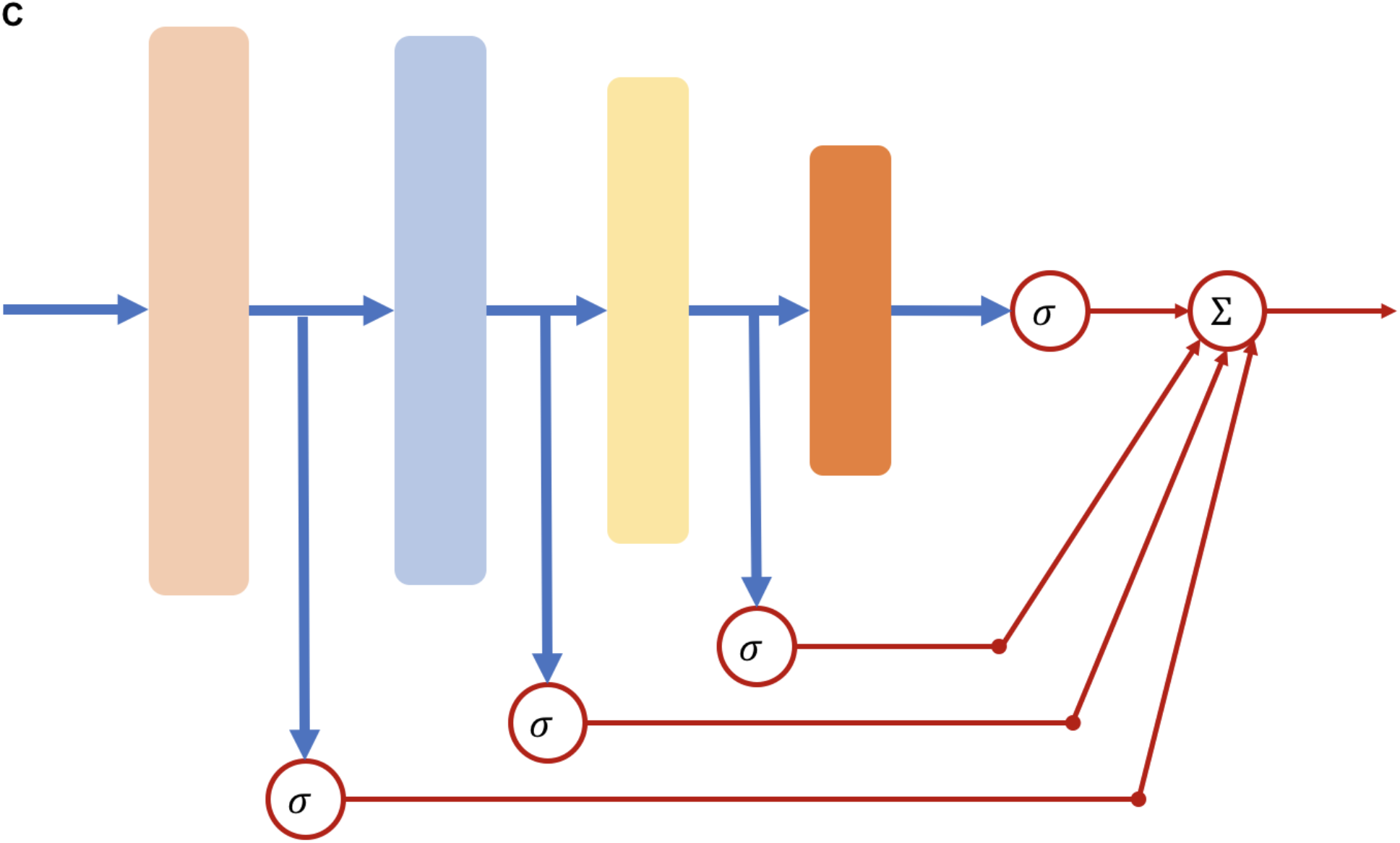
A) The difference between dense and sparse layers. Sparse layers can be arbitrary sparse or patterned sparse. Arbitrary sparse layers are flexible to encode any connection scheme. Patterned sparse layers can make computations more efficient. B) The number of nodes in each layer of the developed P-NET showing a decreasing number of nodes for higher layers. C) A predictive node is connected to each hidden layer in P-NET, and the final prediction is calculated by taking the average of all the predictive elements in the network

**Fig. S2.**
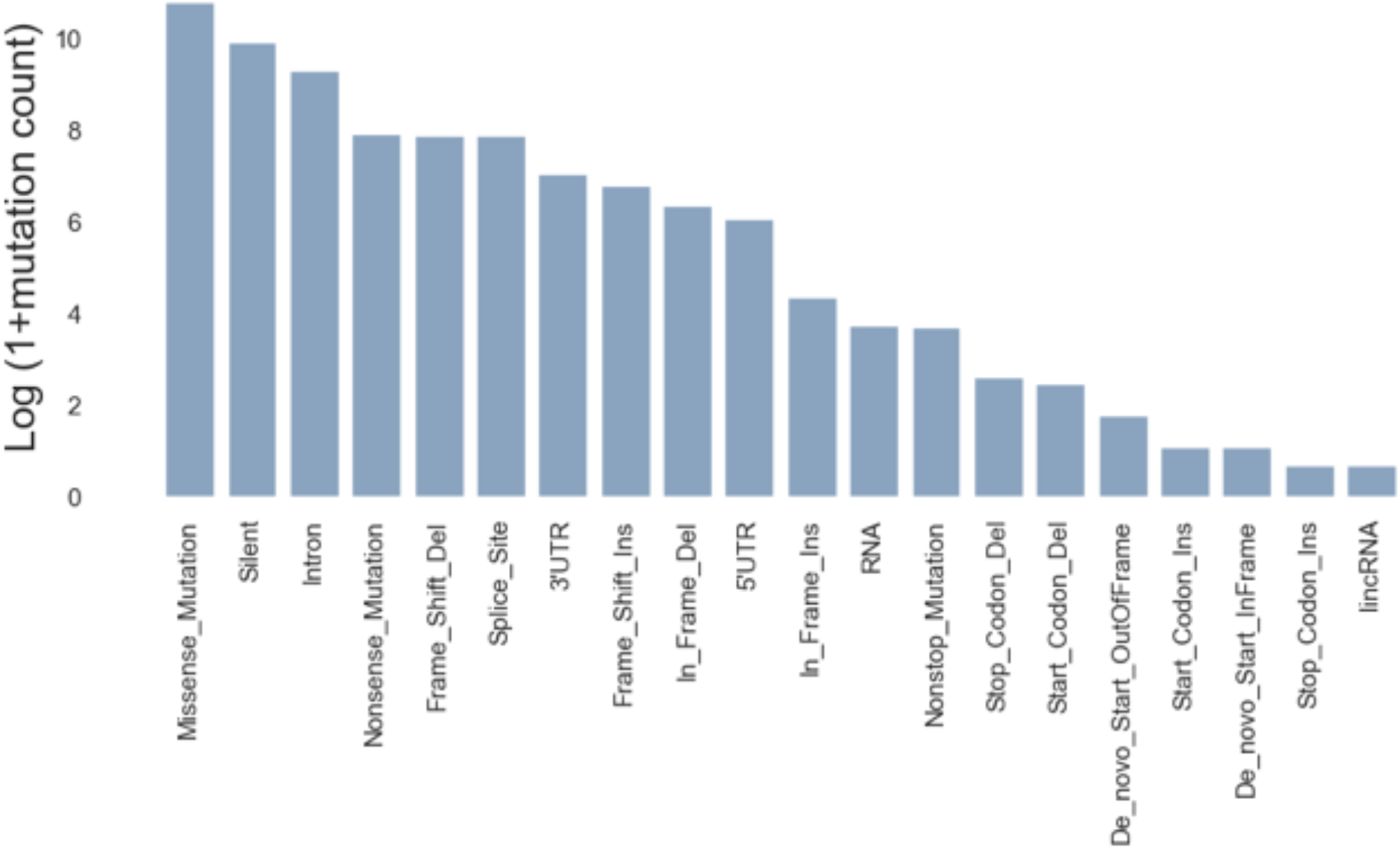
The distribution of mutation types in the P100 dataset. Silent, Intron, 3'UTR, 5'UTR, RNA, and lincRNA mutations are excluded from the modeling.

**Fig. S3.**
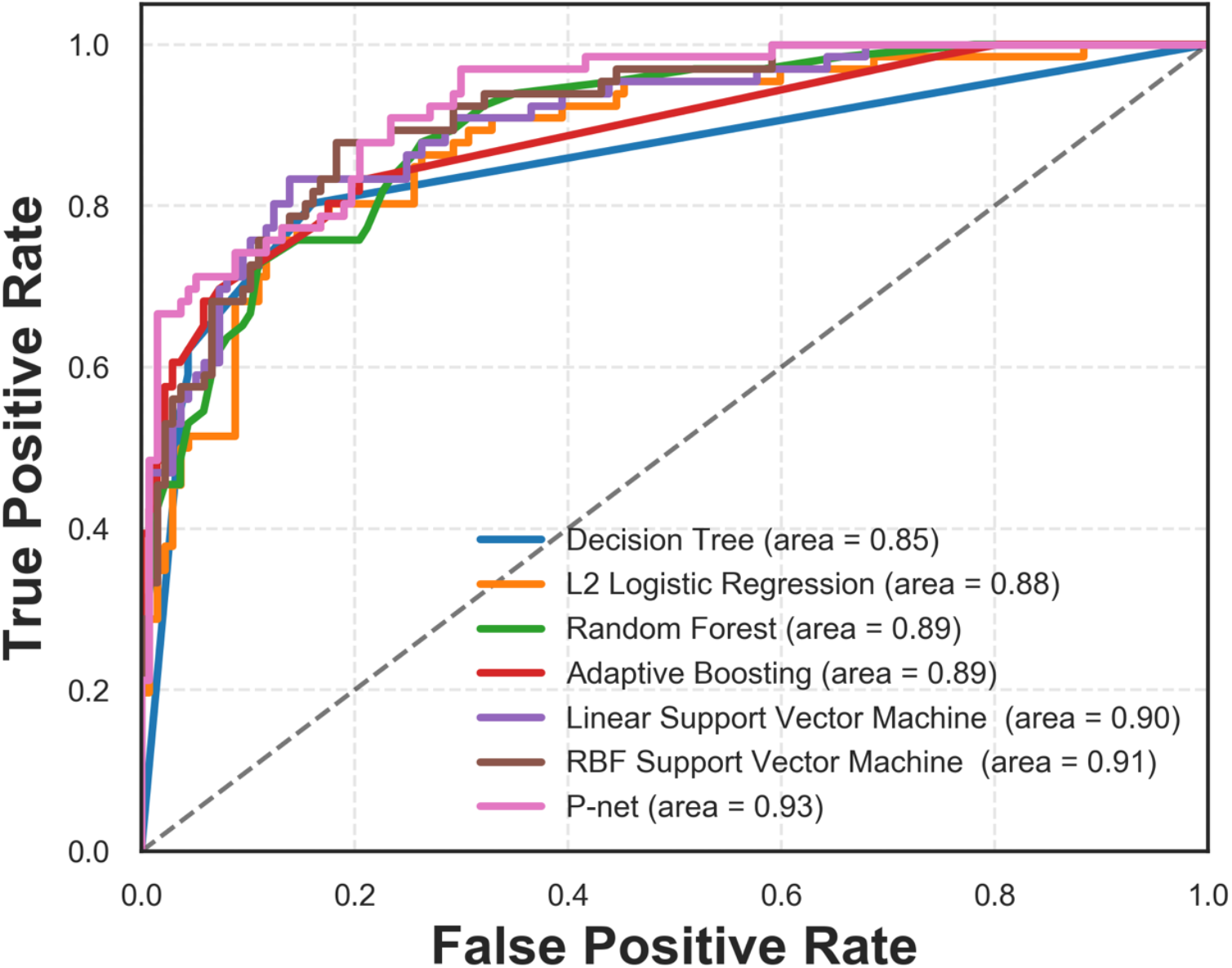
The area under the ROC curve of different models when trained and tested on the P1000 dataset. P-NET outperforms other models leading to higher AUC.

**Fig. S4.**
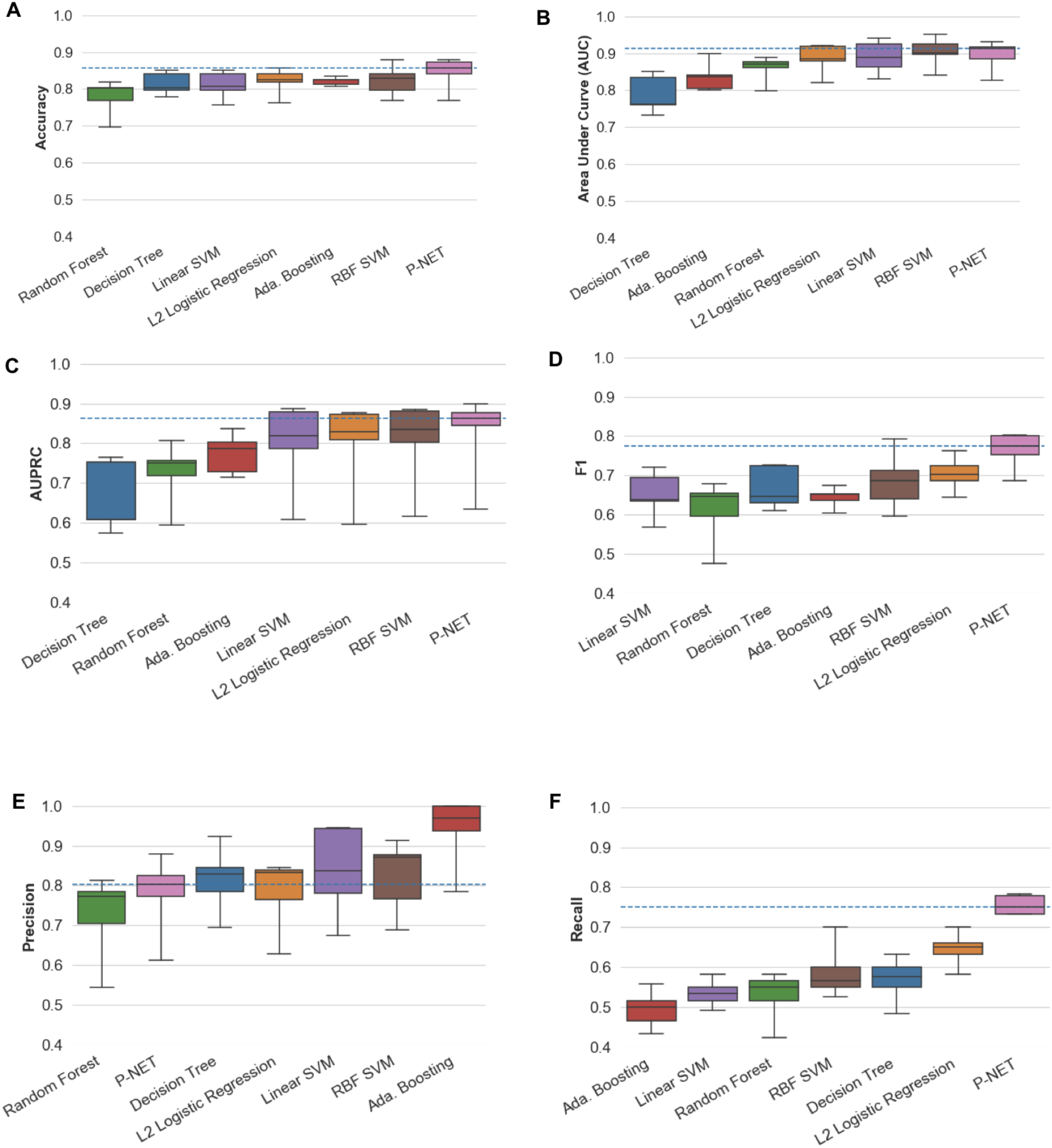
Cross-validation experiment using 5 folds over the development split of the data. P-NET outperforms other models on average using all the metrics (A: Accuracy, B: Area under curve, C: AUPRC, D: F1, and F: recall) except Precision (E).

**Fig. S5.**
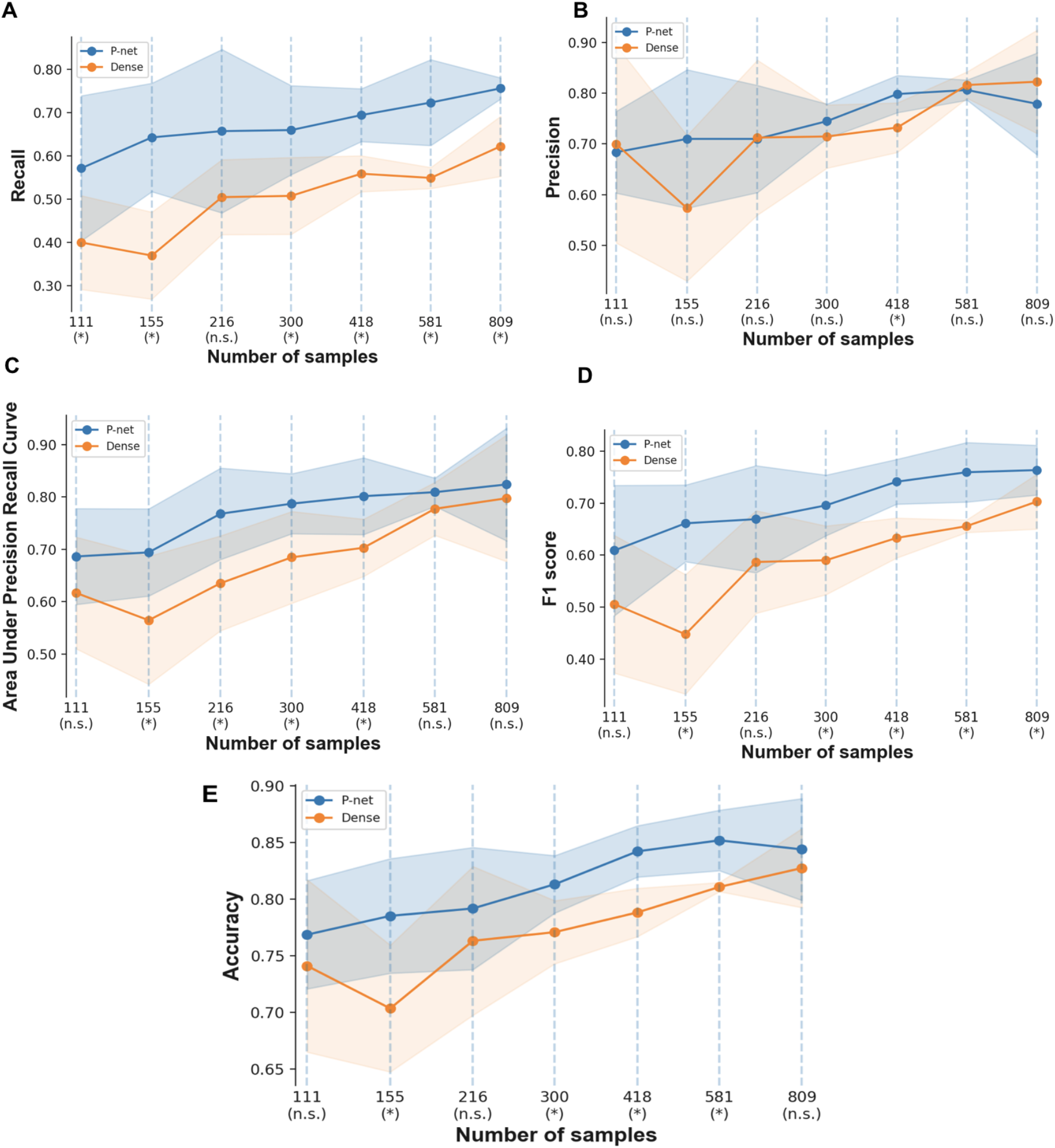
Comparing the performance of P-NET to a dense network with the number of parameters using different sizes of training sets (A: Recall, B: Precision, C: AUPRC, D: F1, E: Accuracy). Sample sizes marked by (*) indicate statistically significant differences while those marked by (n.s.) are not.

**Fig. S6.**
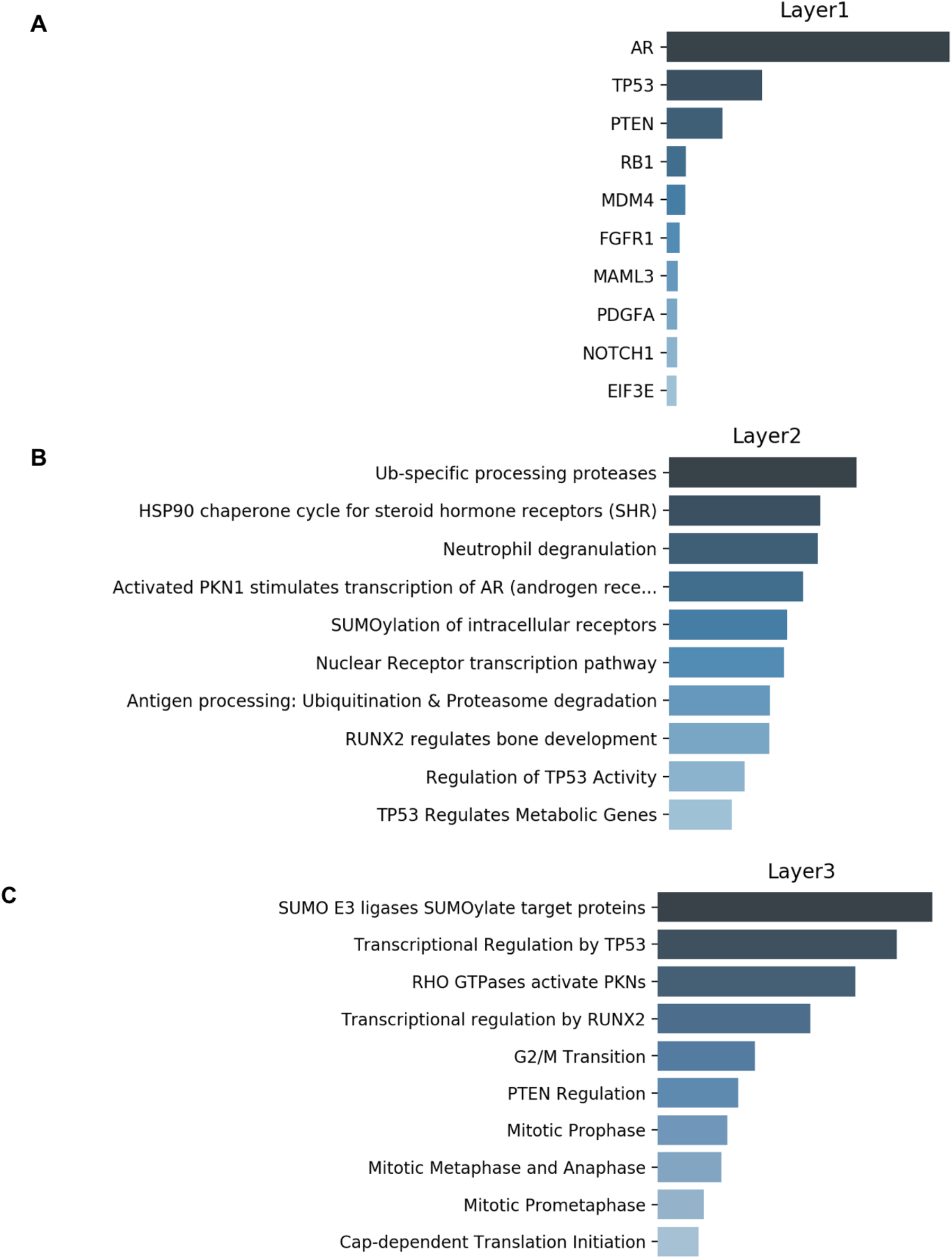
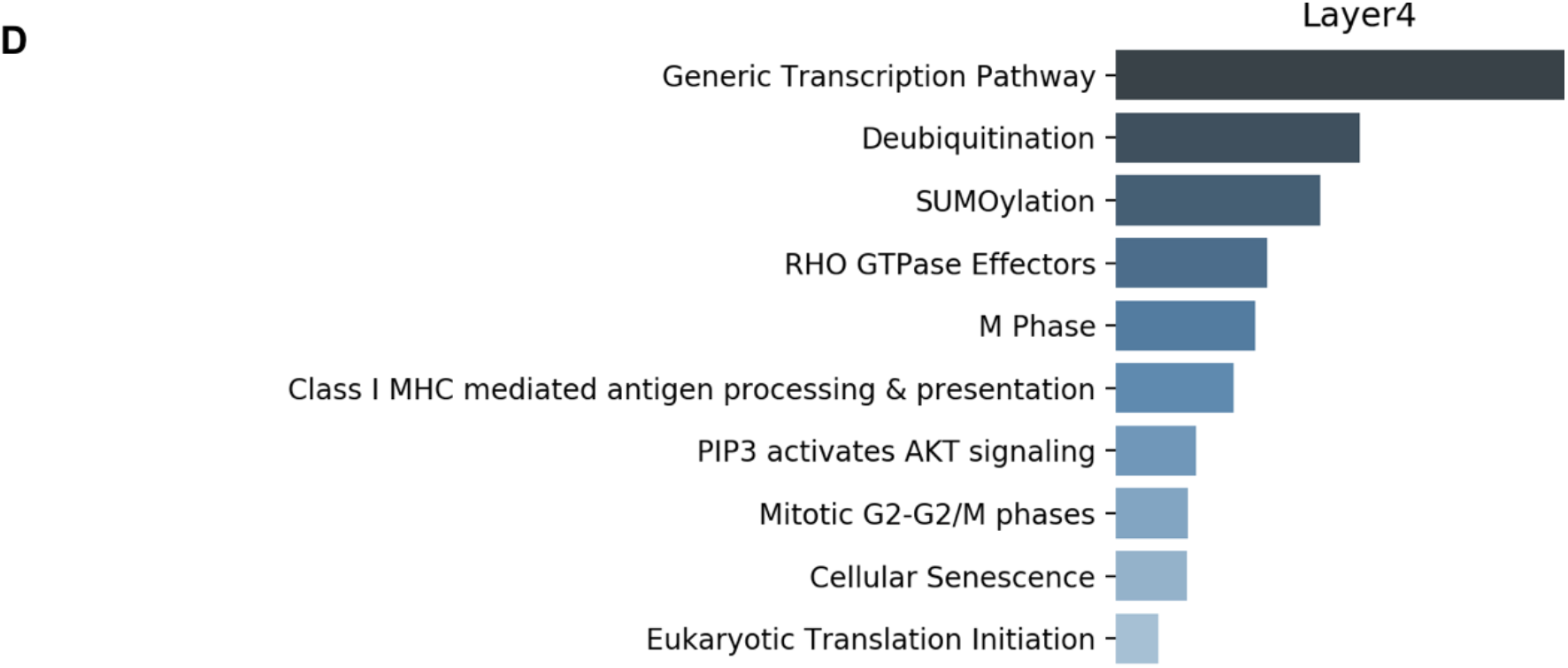
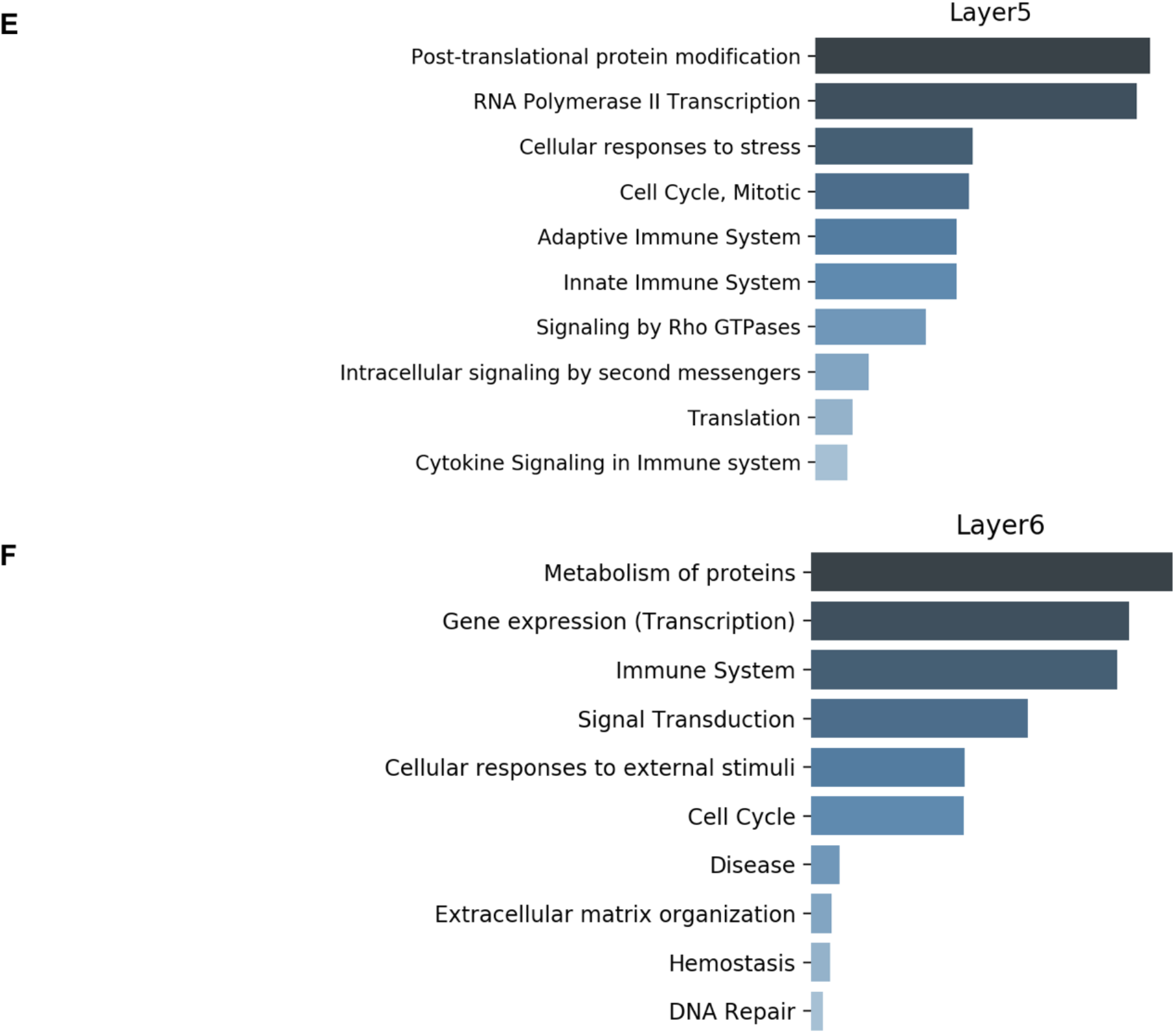
Relative ranking of nodes in each layer

**Fig. S7.**
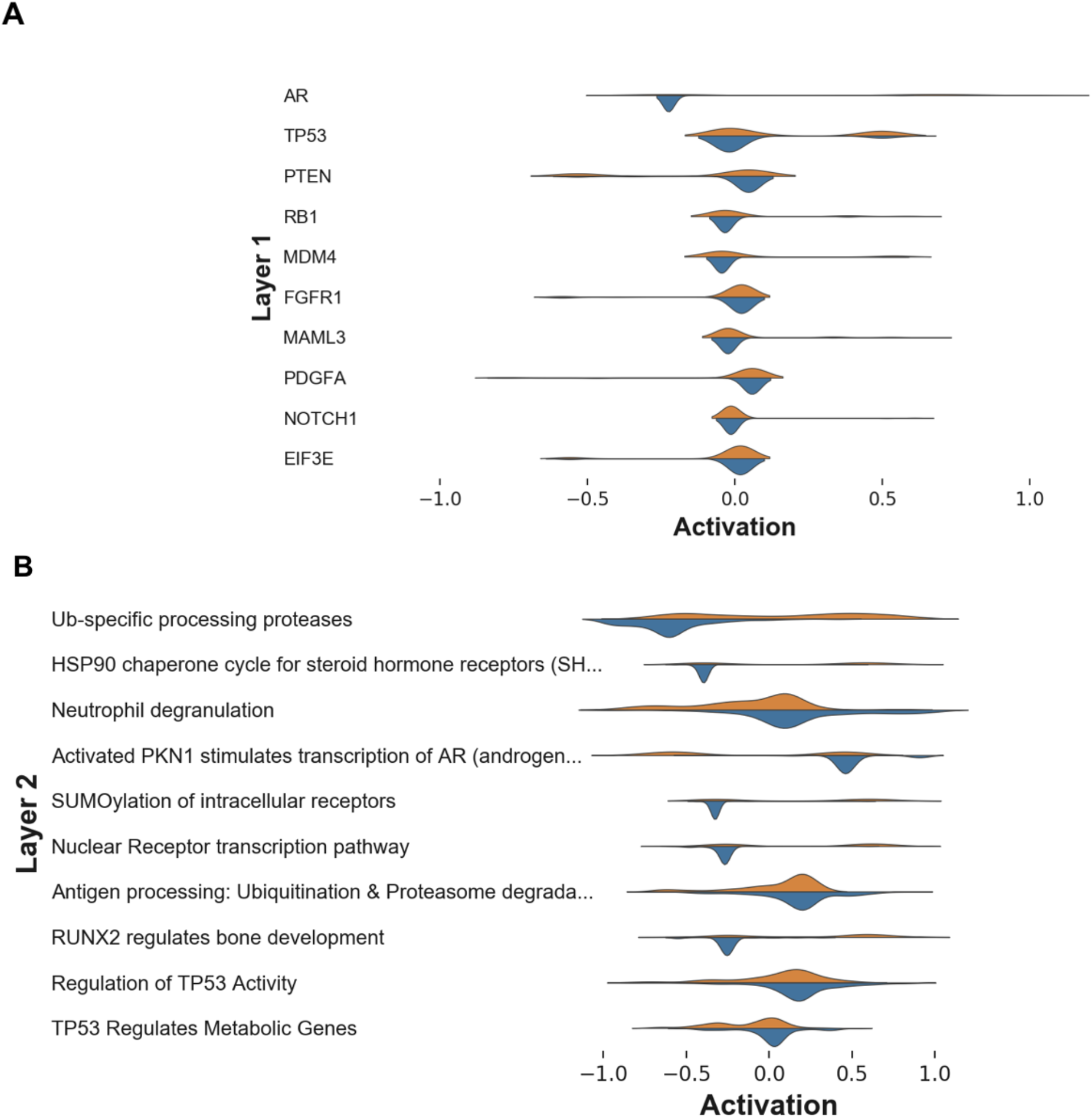
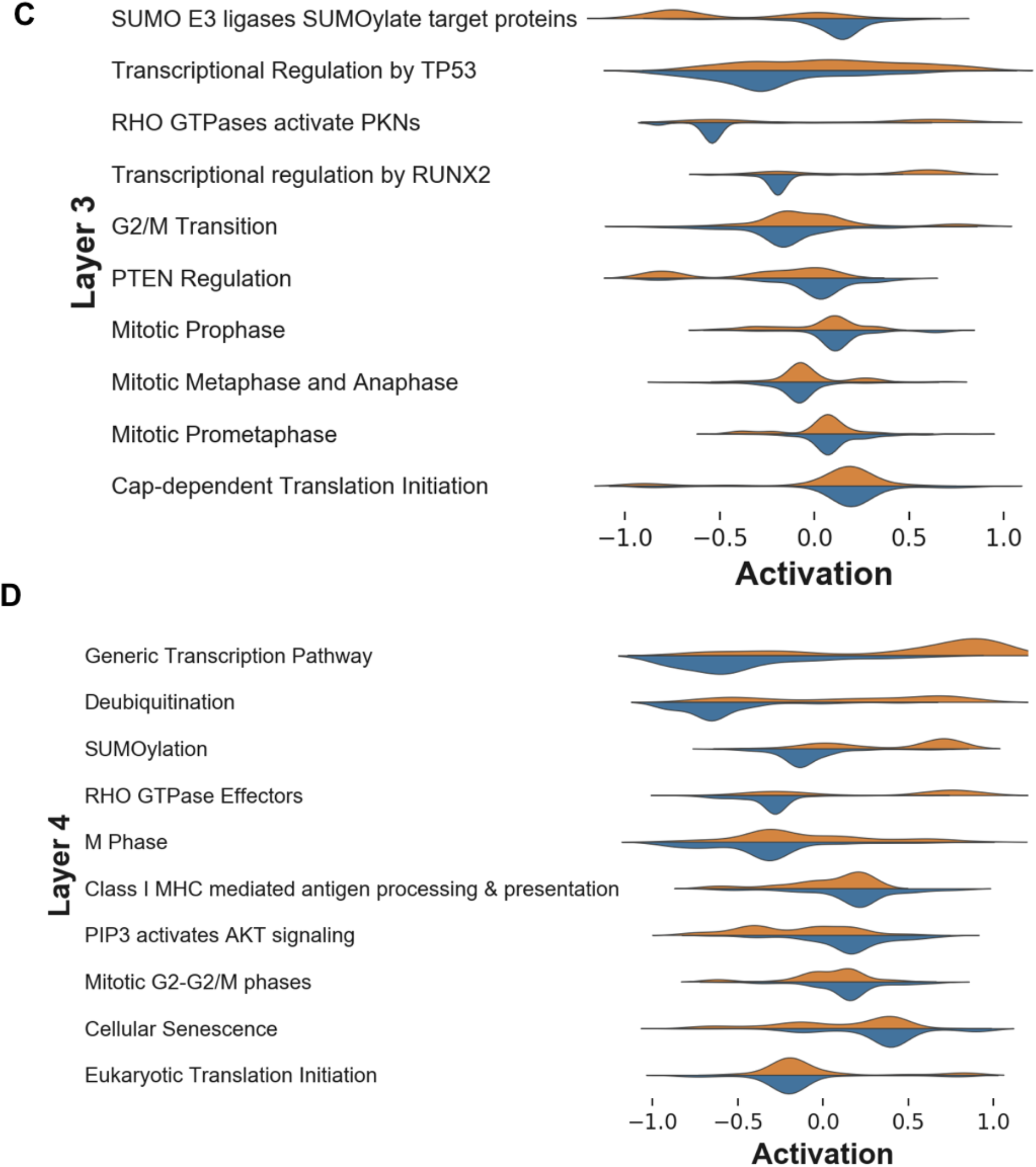
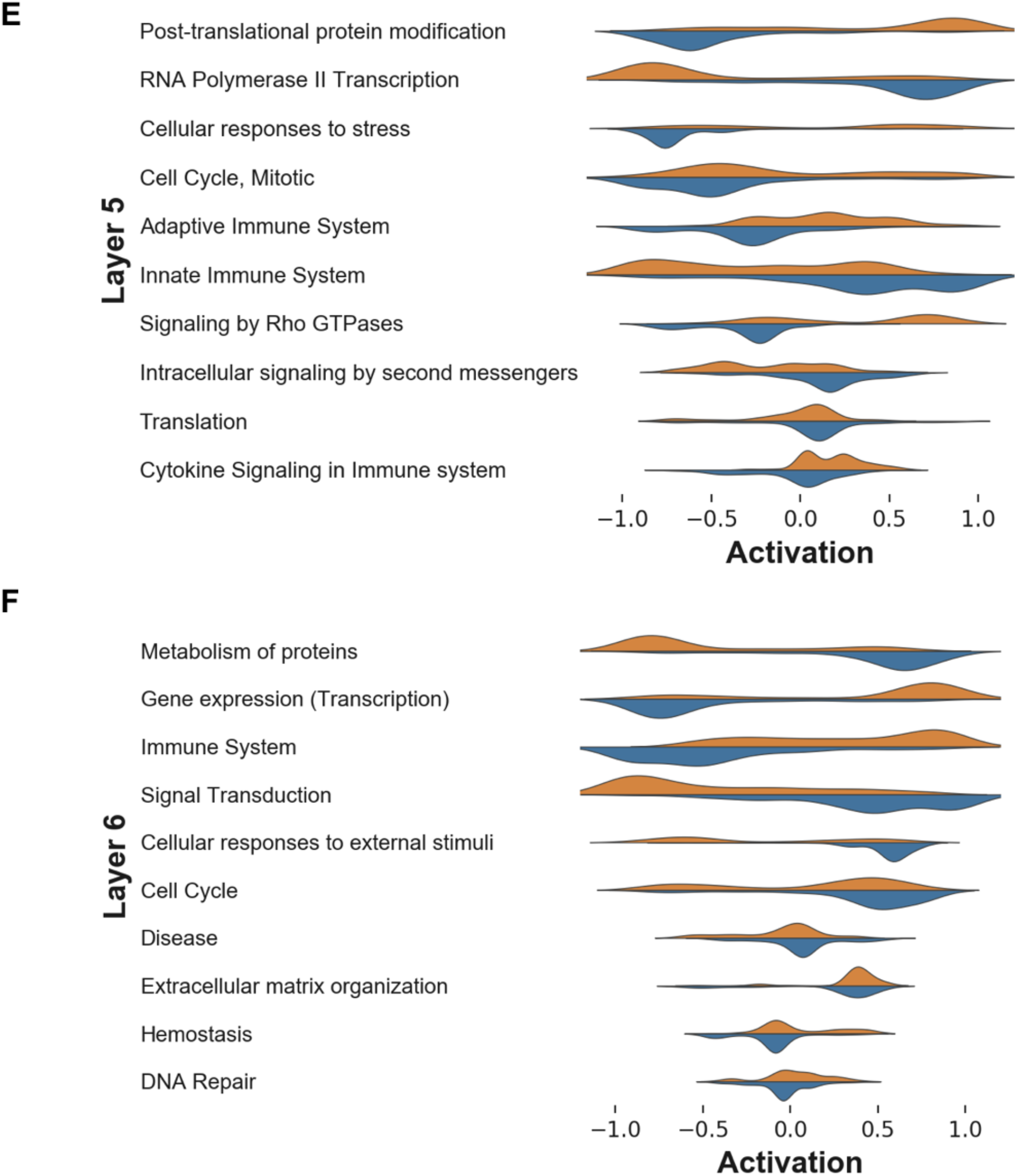
Activation of top ranked nodes in each layer showing better discrimination between sample classes (Primary-blue vs. Metastatic-orange) in higher layers compared lower layers and in top ranked nodes compared to lower ranked ones.

**Fig. S8.**
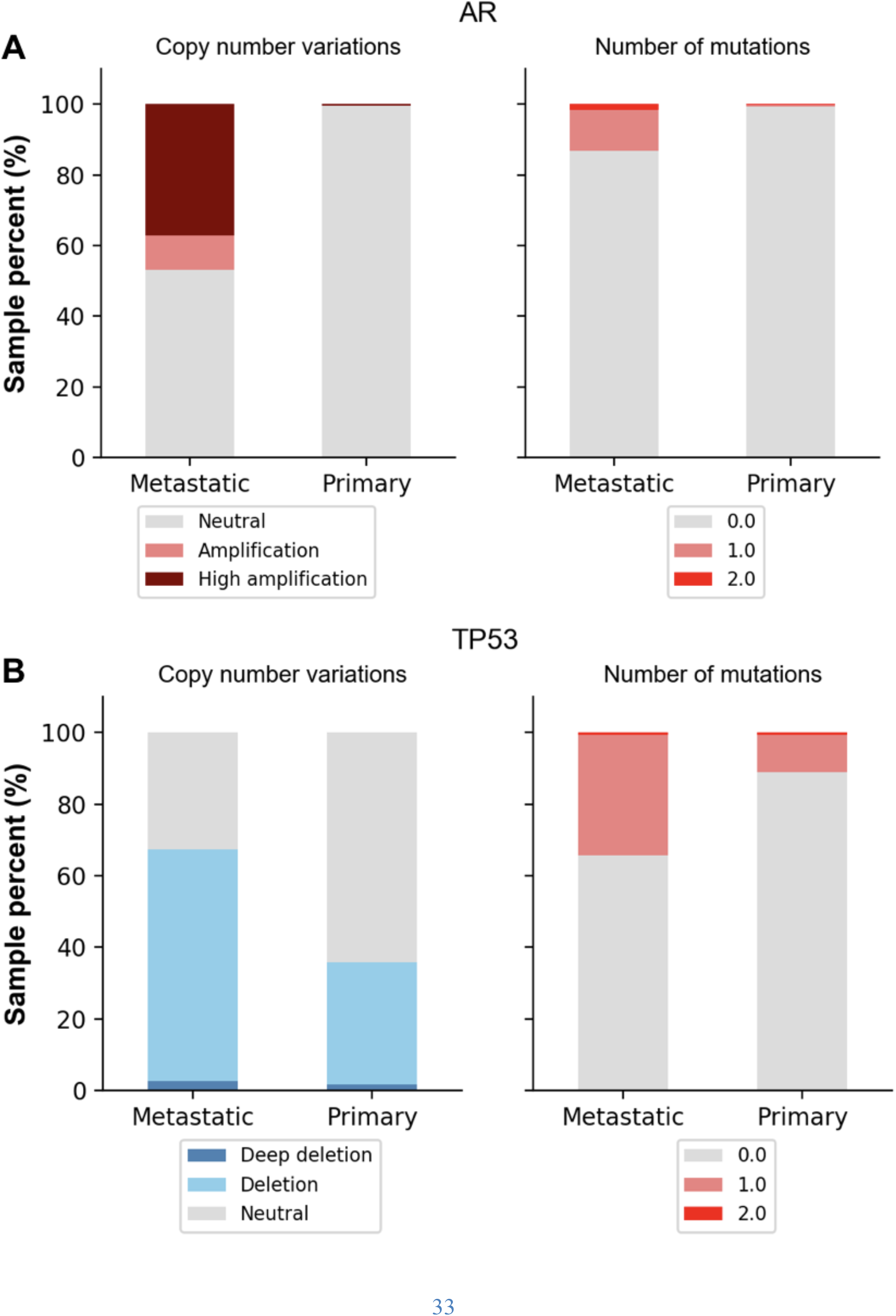
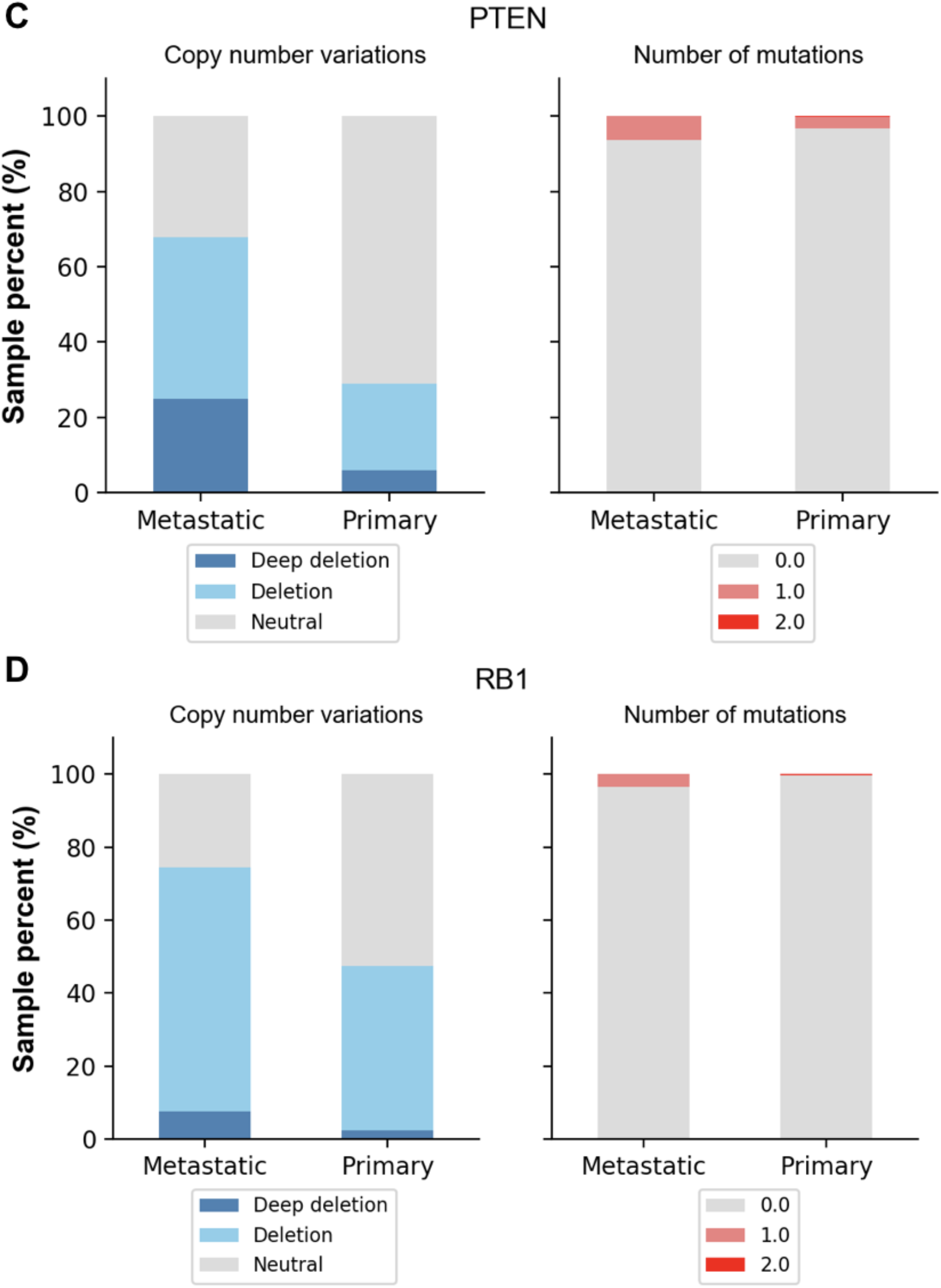
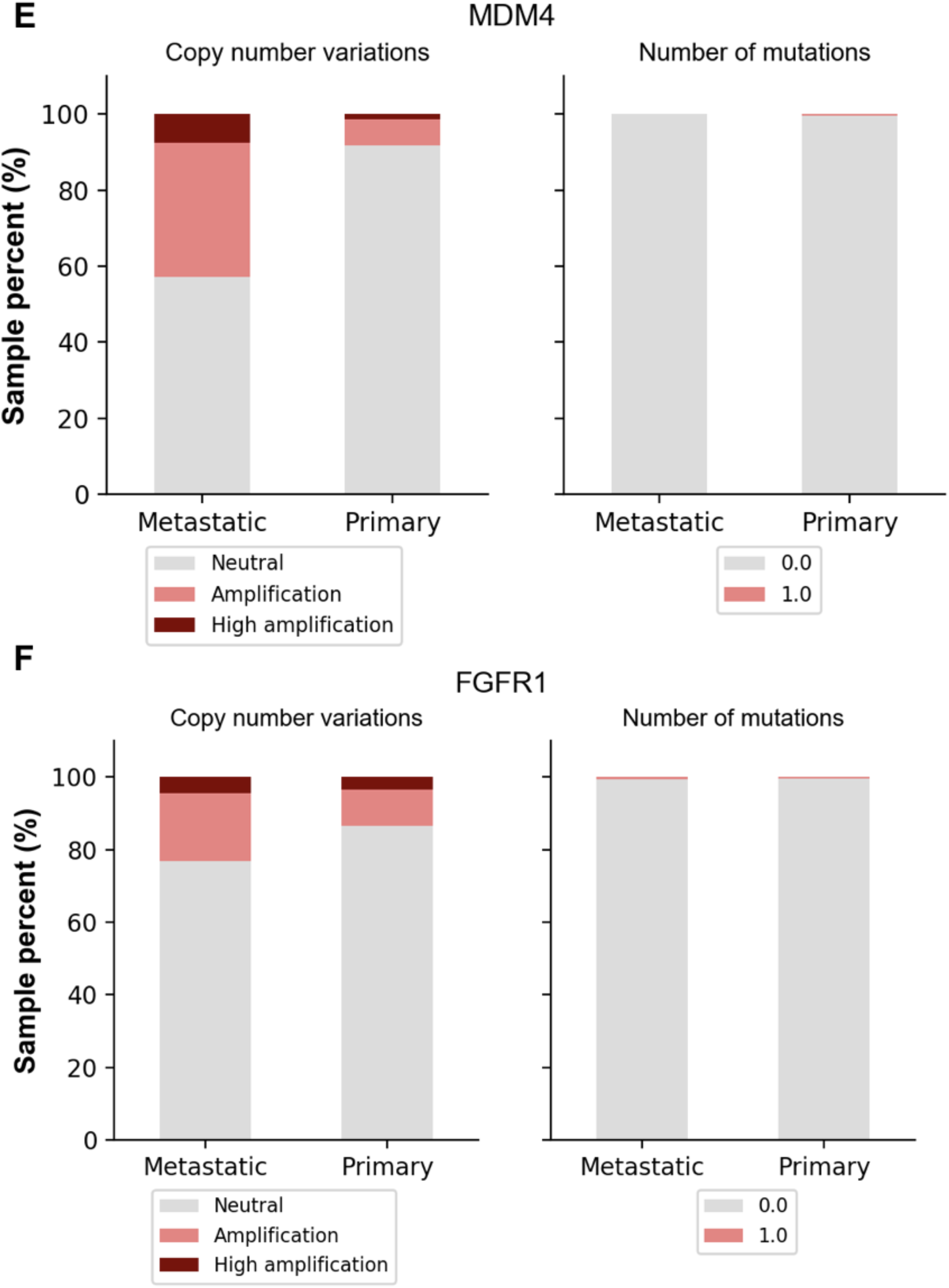
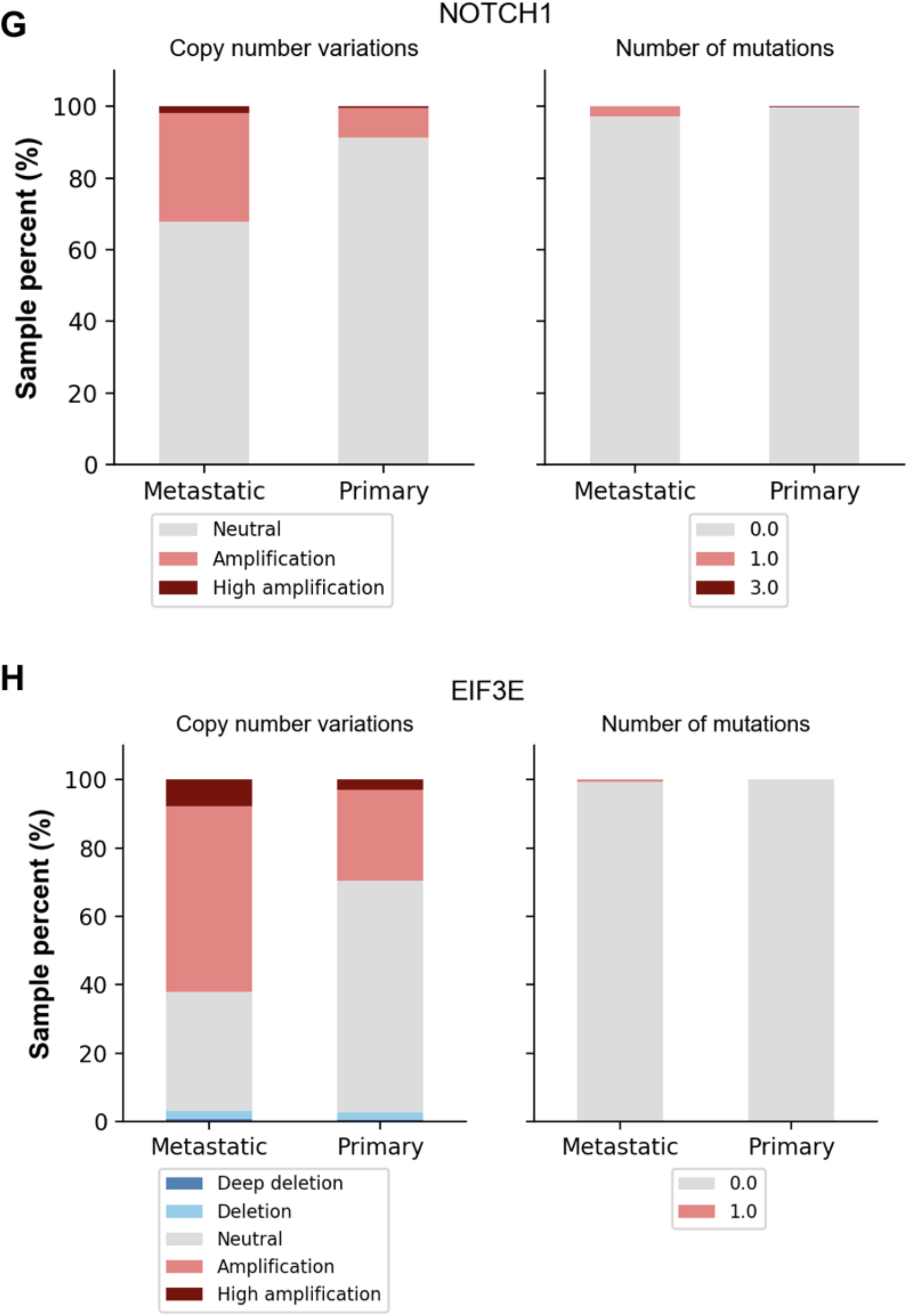
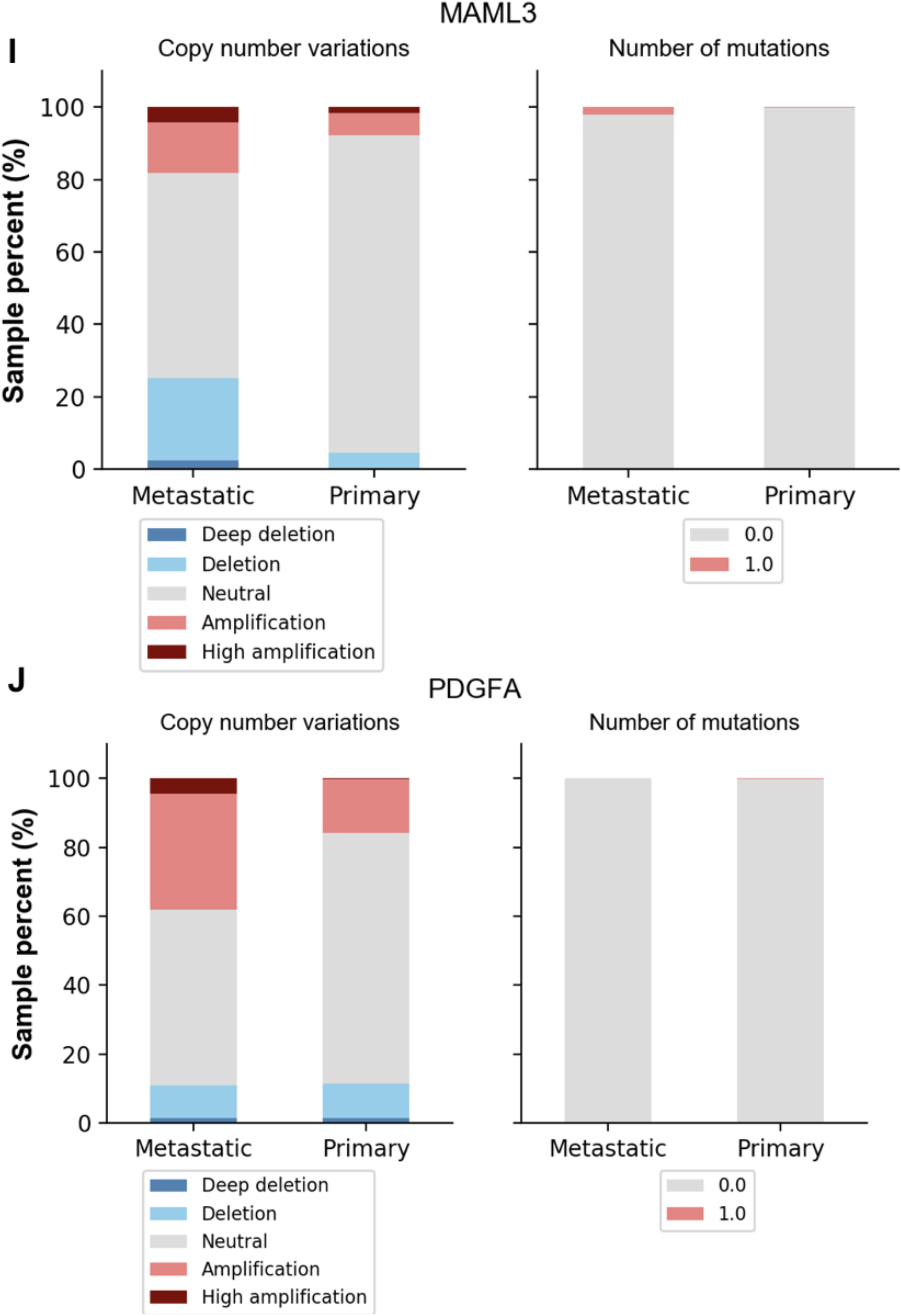
The distribution of mutations and copy number variants of top ranked genes stratified by the class of the samples (Primary vs. Metastatic).

