## Supplementary Data S1 for "Biologically informed deep neural network for prostate cancer classification and discovery"

### Data Links

#### P1000 prostate cancer dataset

Armenia J, Wankowicz SAM, Liu D, Gao J, Kundra R, Reznik E, et al. The long tail of oncogenic drivers in prostate cancer. Nat Genet. 2018;50: 645–651.

<https://static-content.springer.com/esm/art%3A10.1038%2Fs41588-018-0078-z/MediaObjects/41588_2018_78_MOESM6_ESM.xlsx>

<https://static-content.springer.com/esm/art%3A10.1038%2Fs41588-018-0078-z/MediaObjects/41588_2018_78_MOESM4_ESM.txt>

<https://static-content.springer.com/esm/art%3A10.1038%2Fs41588-018-0078-z/MediaObjects/41588_2018_78_MOESM10_ESM.txt>

<https://static-content.springer.com/esm/art%3A10.1038%2Fs41588-018-0078-z/MediaObjects/41588_2018_78_MOESM10_ESM.txt>

<https://static-content.springer.com/esm/art%3A10.1038%2Fs41588-018-0078-z/MediaObjects/41588_2018_78_MOESM5_ESM.xlsx>

External validation dataset

Robinson DR, Wu Y-M, Lonigro RJ, Vats P, Cobain E, Everett J, et al. Integrative clinical genomics of metastatic cancer. Nature. 2017;548: 297–303.

<https://met500.path.med.umich.edu/met500_download_datasets/somatic_v4.csv>

<https://www.dropbox.com/s/62fqw2zgc6ayxvg/Met500_cnv.txt?dl=0>

<https://www.dropbox.com/s/htcx4f09k231l5m/samples.txt?dl=0>

Fraser M, Sabelnykova VY, Yamaguchi TN, Heisler LE, Livingstone J, Huang V, et al. Genomic hallmarks of localized, non-indolent prostate cancer. Nature. 2017;541: 359–364.

<https://static-content.springer.com/esm/art%3A10.1038%2Fnature20788/MediaObjects/41586_2017_BFnature20788_MOESM324_ESM.zip>

<https://static-content.springer.com/esm/art%3A10.1038%2Fnature20788/MediaObjects/41586_2017_BFnature20788_MOESM325_ESM.zip>

Genes

Snapshot of the HUGO gene dataset (accessed 3/21/2018)

<https://www.dropbox.com/sh/kndgiulx7tek9k7/AABZ9XTQ9RfqimI79tTAsbI6a?dl=0>

Pathways

Snapshot of the Reactome pathway dataset (accessed on 09/18/2018)

<https://www.dropbox.com/sh/dhdswefwyh8e9rj/AAD6UmtxFdiJj39_Opc49_Noa?dl=0>
